# Neuregulin 1 type III reduces severity in a mouse model of Congenital Hypomyelinating Neuropathy

**DOI:** 10.1101/386748

**Authors:** Sophie Belin, Francesca Ornaghi, Ghjuvan'Ghjacumu Shackleford, Jie Wang, Cristina Scapin, Camila Lopez-Anido, Nicholas Silvestri, Neil Robertson, Courtney Williamson, Akihiro Ishii, Carla Taveggia, John Svaren, Rashmi Bansal, Schwab H. Markus, Klaus Nave, Pietro Fratta, Yannick Poitelon, Maurizio D’Antonio, M. Laura Feltri, Lawrence Wrabetz

**Author notes:** **Correspondence to:** Lawrence Wrabetz; Hunter James Kelly Research Institute (HJKRI), Jacobs School of Medicine and Biomedical Sciences, University at Buffalo, 701 Ellicott St., Buffalo, NY, 14203, USA.

## Abstract

Myelin sheath thickness is precisely regulated and essential for rapid propagation of action potentials along myelinated axons. In the peripheral nervous system, extrinsic signals from the axonal protein neuregulin 1 type III regulate Schwann cell fate and myelination. Here we ask if modulating neuregulin 1 type III levels in neurons would restore myelination in a model of congenital hypomyelinating neuropathy (CHN). Using a mouse model of CHN, we rescued the myelination defects by early overexpression of neuregulin 1 type III. Surprisingly, the rescue was independent from the upregulation of *Egr2* or essential myelin genes. Rather, we observed the activation of MAPK/ERK and other myelin genes such as peripheral myelin protein 2 (*Pmp2*) and oligodendrocyte myelin glycoprotein (*Omg*). We also confirmed that the permanent activation of MAPK/ERK in Schwann cells has detrimental effects on myelination. Our findings demonstrate that the modulation of axon-to-glial neuregulin 1 type III signaling has beneficial effects and restores myelination defects during development in a model of CHN.

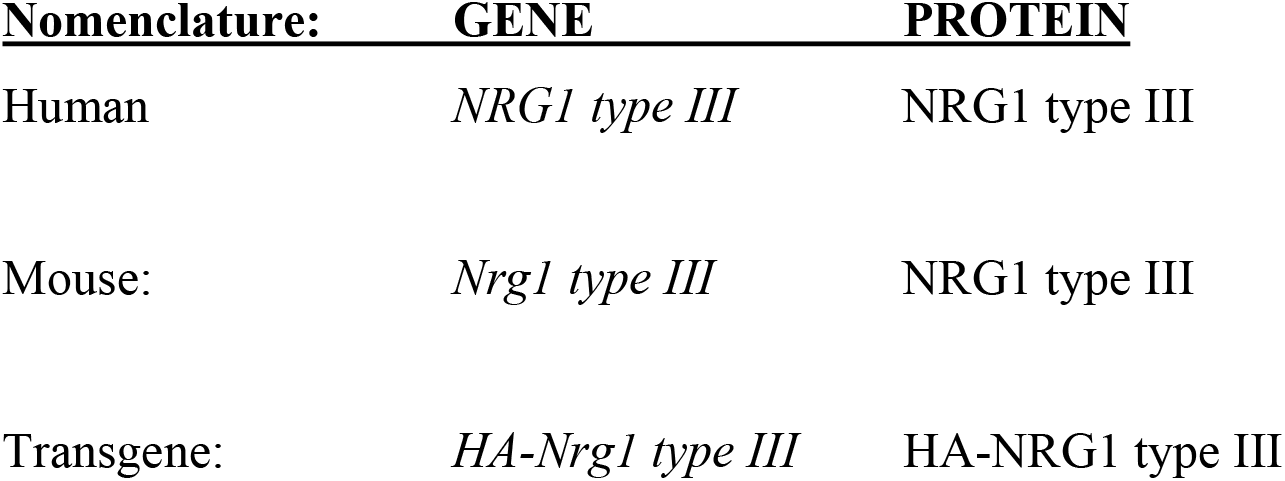

## Introduction

The neuregulin 1 (NRG1) family of ligands and cognate ErbB tyrosine kinase receptors (heterodimeric ErbB2/B3), regulate much of Schwann cell development (Birchmeier and Nave, 2008; Garratt et al., 2000; Lin et al., 2000; Meyer and Birchmeier, 1995; Michailov et al., 2004; Nave and Salzer, 2006; Riethmacher et al., 1997; Taveggia et al., 2005). NRG1 function depends on isoform and amount of expression (Fricker et al., 2011; Gomez-Sanchez et al., 2009; Stassart et al., 2013). In particular, the level of expression of the axonal membrane bound NRG1 type III is a potent instructive signal that determines whether a Schwann cell will form myelin, as well as its thickness (Michailov et al., 2004; Taveggia et al., 2005). A series of mouse genetics experiments have elucidated the complex molecular network downstream of NRG1/ErbB2/ErbB3 that regulates myelination during development and remyelination (Fledrich et al., 2014; Pereira et al., 2012; Stassart et al., 2013). Intrinsic Schwann cell signals downstream of NRG1 include PI3K/AKT, MAPK/ERK and calcineurin/NFATc4, which are required to precisely regulate myelin gene transcription and protein synthesis required for myelin formation (Brinkmann et al., 2008; Grossmann et al., 2009; He et al., 2010; Kao et al., 2009; Maurel and Salzer, 2000; Newbern et al., 2011; Ogata et al., 2004).

Expressed almost exclusively by myelin-forming Schwann cells, myelin protein zero (MPZ/P0) is the major protein in myelin of the peripheral nervous system (PNS) and is necessary for normal myelin structure and function (Giese et al., 1992). In addition to structural interactions with the membrane, the P0 cytoplasmic tail is thought to contain motifs that assemble signal complexes to stimulate myelin gene expression and therefore modulate myelin thickness (Giese et al., 1992; Xu et al., 2000). A human nonsense mutation in the *Mpz* gene generates a premature stop codon in the P0 protein at glutamine 215 (P0Q215X). This truncates the P0 cytoplasmic tail and causes a severe congenital hypomyelinating neuropathy (CHN) in patients, characterized by weakness, hypotonia and areflexia from birth, associated with a severely reduced nerve conduction velocity (Mandich et al., 1999; Warner et al., 1996). The molecular mechanisms underlying how truncation of P0 triggers hypomyelination are not well understood.

In this study, we used a mouse model of CHN with the targeted mutation of *Mpz* encoding the P0Q215X mutant protein. Mice heterozygous for Q215X (*Mpz*^Q2l5XI/+^) manifest diminished motor performance, and hypomyelination that recapitulates a mild Q215X human nerve pathology (Fratta et al., 2018). Genetic experiments suggest that the Q215X phenotype results from a combination of loss and gain-of function. Here we find that the neuregulin axis is intact in Q215X nerves, but myelin gene expression is reduced. Although overexpression of the *Mpz* gene does not impact on the Q215X phenotype (Fratta et al., 2018), we reasoned that augmented activity of *Nrg1 type III* might compensate for hypomyelination in CHN by increasing myelin gene expression and myelin thickness (Michailov et al., 2004). To test this, we crossed Q215X mice with a transgenic mouse overexpressing *HA-Nrg1 type III* in neurons (Velanac et al., 2012). We observed an improvement in myelin thickness and nerve conduction velocity (NCV) in *Nrg1 type III*:Q215X mice. Transcriptional and biochemical analyses identified MAPK/ERK as the likely signaling pathway responsible for rescuing Q215X defects in Schwann cells.

## Results

### Myelin protein levels are reduced in Q215X mice

We showed that *Mpz*^Q215X/+^ mice present an hypomyelination in sciatic nerves (Fig. 1A, B), caused by a technical loss of function and a dose-dependent gain of abnormal function (Fratta et al., 2018). Thus, although the Q215X mutant allele caused a mild phenotype, the Q215X model displayed features of a human CHN with non-progressive hypomyelination throughout development (Fratta et al., 2018). To determine if the hypomyelination in *Mpz^Q215X/+^* Schwann cells is caused by a misregulation of myelin genes, we quantified RNA and protein levels of the major myelin genes in Q215X mutants during development. Both Q215X heterozygote and homozygote sciatic nerves showed a reduction of *Pmp22* and *Mag* mRNA at early time points P5 and/or P10, when myelination is initiating (Fig. 1 C-F). In addition, Q215X heterozygote and homozygote sciatic nerves had reduced levels of PMP22, MAG and MBP at P5 (Fig. 1 G-J and Fig. EV1). We analyzed the mRNA and protein levels for *Egr2*, a key transcriptional regulator of myelin genes, but did not find any dysregulation, suggesting that the underexpression of myelin genes in Q215X is independent of *Egr2* (Fig. 1 F, J). These results suggest that the Q215X allele affect myelin protein expression and is well suited for the study of therapeutical approaches aiming at rescuing hypomyelination.

**Figure 1.**
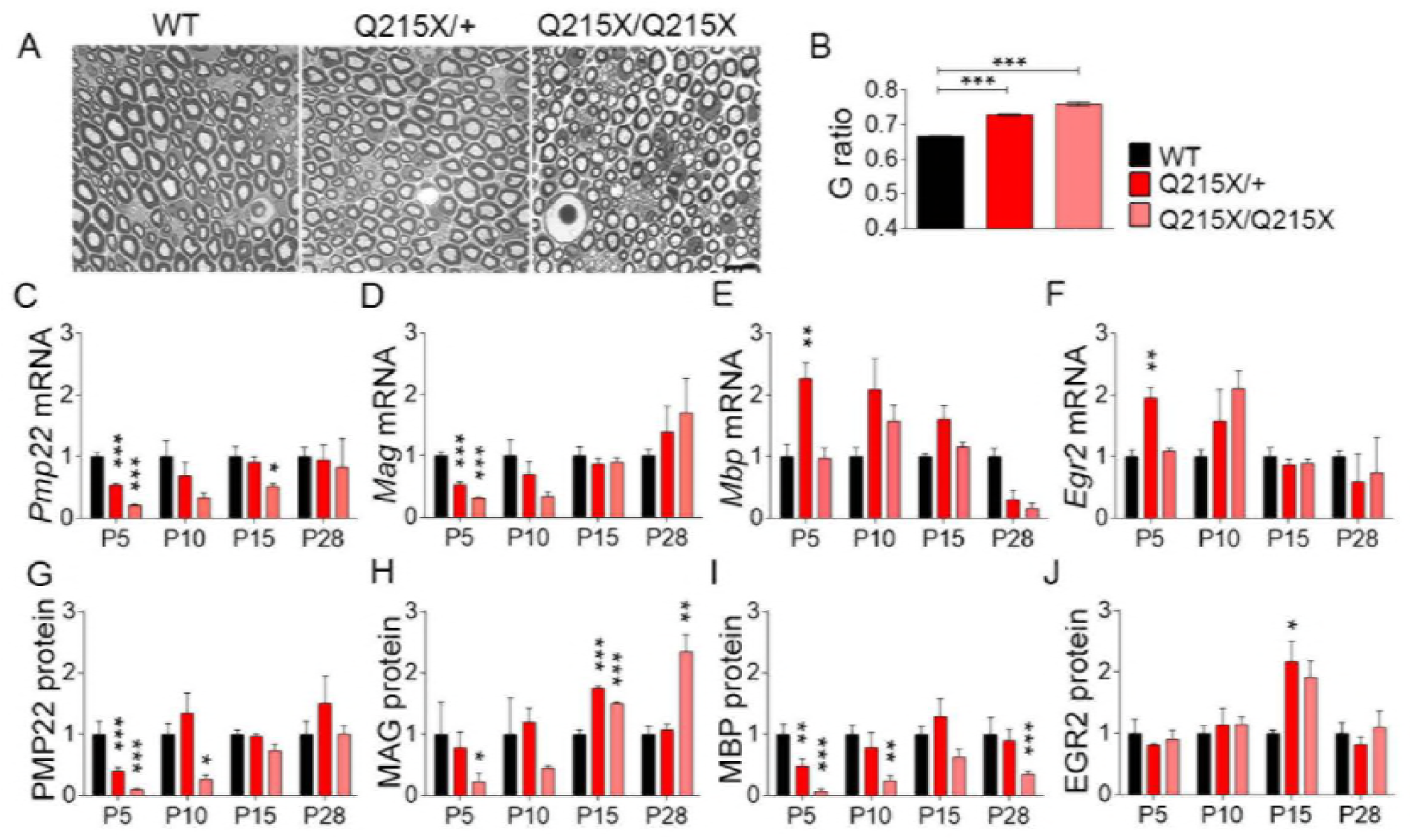
The Q215X mutation in P0 causes hypomyelination and alters myelin genes expression. (A) Semithin cross-sections stained with toluidine blue from wildtype (WT), P0Q215X heterozygote (Q215X/+) and P0Q215X homozygote (Q215X/Q215X) sciatic nerves at 1 month. Q215X/+ and Q215X/Q215X fibers are hypomyelinated as compared to WT. Scale bars, 10 μm. (**B**) g-ratio was increased in Q215X mutants from 0.66±0.0005 (WT) to 0.72±0.003 (Q215X/+) and 0.76±0.005 (Q215X/Q215X). (**C-F**) mRNA relative quantification by real time PCR for *Pmp22, Mag, Mbp* and *Erg2* in WT, Q215X/+, Q215X/Q215X sciatic nerves at P5, P10, P15 and P30. (**G-J**) Densitometries from western blot analyses of PMP22, MAG, MBP and EGR2. Western blot analyses were performed on sciatic nerves lysates of WT, Q215X/+, Q215X/Q215X at P5, P10, P15 and P30. Representative blots are shown in Fig. EV1. Analyses were performed using three mice per genotype. Error bars indicate s.e.m. Statistical analyses were performed using one-way ANOVA with Bonferroni’s multiple comparison. * *P* < 0.05, ** *P* < 0.01, *** *P* < 0.001.

### Overexpression of NRG1 type III restores myelin thickness in Q215X mice

NRG1 type III expressed on axons determines whether Schwann cells will form myelin, and its thickness, through the activation of the master myelin gene transcription factor *Egr2* (Kao et al., 2009; Taveggia et al., 2005). Overexpressing NRG1 type III isoform in sensory and motor neurons significantly increases myelin thickness in the peripheral nervous system (Michailov et al., 2004 Velanac et al., 2012).

Q215X mutants present a hypomyelination with decreased myelin gene expression, likely due to a loss of transcriptional signals. Thus, we expect that introducing exogenous expression of HA-NRG1 type III in a hypomyelinating context (Q215X) could activate transcription of myelin genes and restore myelin thickness. We bred Q215X mice with the HA-NRG1 type III hemizygous transgenic mouse model (Nrg1t3) that overexpresses HA-NRG1 type III under the control of the neuronal Thy1.2 promoter (Campsall et al., 2002; Caroni, 1997).

Morphological and ultrastructure analysis of Nrg1t3;Q215X sciatic nerves were performed at one month of age and compared to Q215X heterozygote (Q215X) littermates. First, we confirmed by morphologic measurements that the newly generated Q215X and Nrg1t3 mice present hypomyelination and hypermyelination, with respectively 0.72±0.01 and 0.53±0.02 as G ratio, as compared to 0.67±0.005 in wildtype littermates (Fig. 2 A-C). Second, we observed that double mutant animals present a dramatic recovery of myelin thickness when compared to Q215X, and return to a myelin thickness close to wild-type (WT) animals with a G ratio of 0.62±0.01 (Fig. 2A-C).

As previously described with transgenic expression of HA-NGR1 type III (Michailov et al., 2004; Velanac et al., 2012), the increase in myelin thickness was predominantly observed in small caliber axons (Fig. 2C). We also observed that Nrg1t3 animals present increased internodal length as well as more myelinated small caliber fibers from 1 to 2 μm (Fig. EV2A-C).

As expected, in Q215X animals the hypomyelination observed in one-month old animals correlated with a reduction of nerve conduction velocities (Fig. 2D). In the Nrg1t3 mice sciatic nerve axons were well myelinated, and there was no alteration of nerve conduction velocity and F wave latency (Fig. 2D, E). The increase of myelin thickness in Nrg1t3;Q215X led to an improvement of nerve conduction velocities (Fig. 2D). However, the nerve conduction velocities do not return to the wild-type level, likely due to the pronounced effect of Nrg1 type III on small caliber fibers that contributes in a minor way to electrophysiological measurements of nerve conduction velocity. The F-wave latencies were also improved in Nrg1t3;Q215X, suggesting an increase in proximal myelination (Fig. 2E).

P0 is essential for proper myelin compaction (Ding and Brunden, 1994; Kirschner and Ganser, 1980; Lemke, 1988; Shapiro et al., 1996). To exclude that the changes in myelin thickness were due to a defect in compaction, we analyzed the periodicity of myelin sheath, but no difference between mutants and WT were found (Fig. 2F, G).

**Figure 2.**
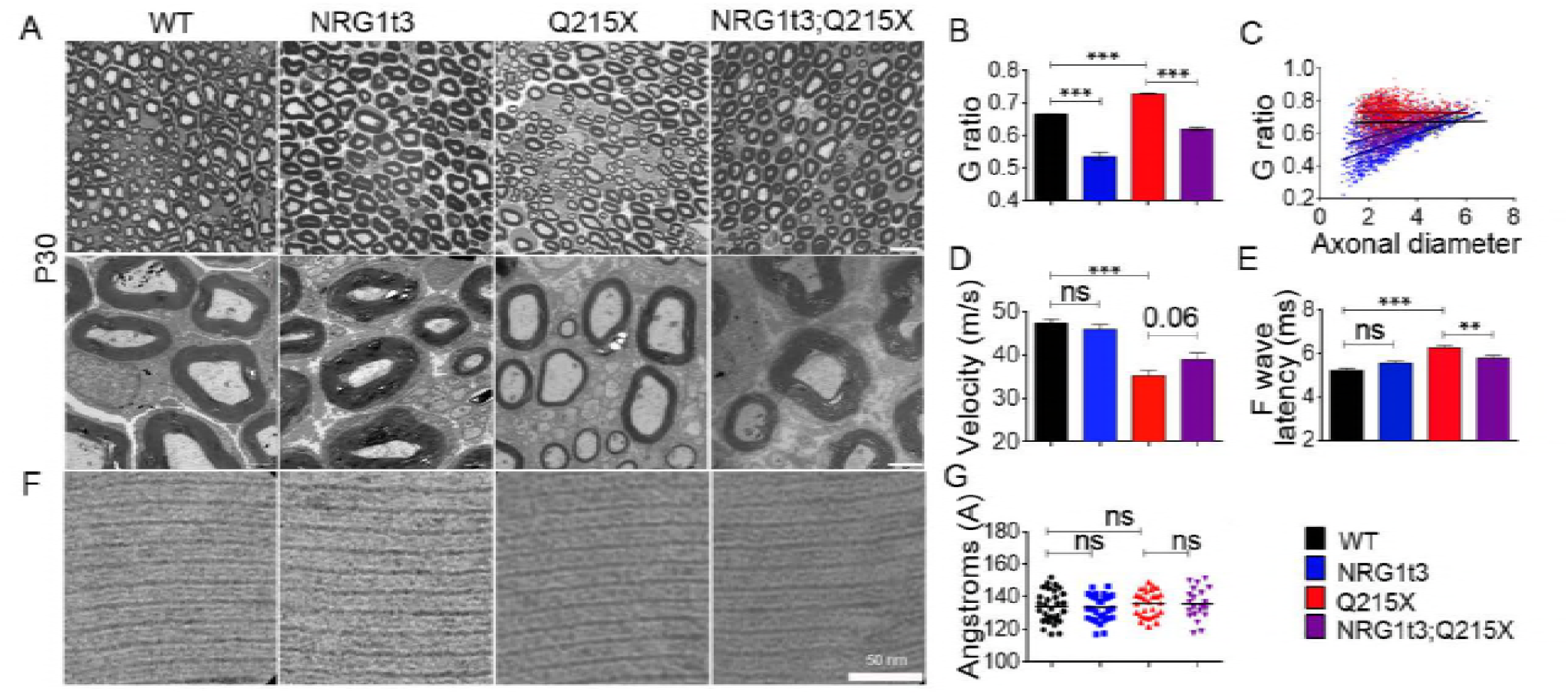
Neuronal overexpression of NRG1 typeIII restores myelination in P0Q215X. (**A**) Semithin cross-sections stained with toluidine blue (top panel) and electron micrographs (bottom panel) from wildtype (WT), Nrg1t3 hemizygote (Nrg1t3), P0Q215X heterozygote (Q215X) and Nrg1t3 hemizygote and P0Q215X heterozygote (Nrg1t3;Q215X) sciatic nerves at one month. NRG1t3 and Nrg1t3;Q215X fibers are hypermyelinated as compared to WT and Q215X. Scale bars, 10 μm (top panel) and 2 μm (bottom panel). (**B**) g-ratio was increased in Q215X (0.72±0.003) and decreased in Nrg1t3 (0.53±0.02), indicating hypomyelination and hypermyelination, respectively. (**C**) g-ratio as a function of axonal diameter showed preferential hypermyelination of small fibers in Nrg1t3 and Nrg1t3;Q215X as compared to WT and Q215X. (**D**) Neurophysiological studies showed a significant reduction of nerve conduction velocity in Q215X mice when compared to WT (47.28±0.97 m/s). A recovery of nerve conduction velocities can be observed in Nrg1t3;Q215X (39±1.5 m/s) as compared to Q215X (35±1.28 m/s) (*P* = 0.06). (**E**) F wave latencies were increased in Q215X (6.54±0.11 m/s) when compared to WT (5.32±0.08 m/s). A significant recovery of F wave latencies is observed in Nrg1t3;Q215X (6±0.12 m/s), suggesting a proximal myelination improvement. (**F-G**) Myelin compaction is not altered in single or double mutant as indicated in electron micrographs (**F**) and periodicity quantification (**G**). Scale bar, 50 nm. Analyses were performed using three mice per genotype. Error bars indicate s.e.m. Statistical analyses from three independent experiments using oneway ANOVA with Bonferroni’s multiple comparison test. ** *P* < 0.01, *** *P* < 0.001.

Q215X mutant displayed early postnatal altered motor performance (Fratta et al., 2018). Therefore, we assessed the effect of HA-NRG1 type III on motor performance using the grid-walking test at P15. Unlike the first transgenic mouse model generated which developed mild tremor (Michailov et al., 2004), no visible tremor phenotype was described in the Nrg1t3 transgenic mouse (Velanac et al., 2012). However, the grid-walking test at P15 revealed motor coordination deficits (more ‘foot faults’ were recorded as compared to WT) in Nrg1t3 mice. This interfered with the behavioral investigations to assess a possible motor rescue in Nrg1t3;Q215X mice (Appendix Fig. S1). Multiple studies in human and rodents link mutations in *NRG1* gene or imbalanced ErbB4/NRG1 signaling to schizophrenia-like behaviors accompanied by increased anxiety levels and disrupted sensorimotor gating, which were recently associated to an impaired synaptic plasticity (Agarwal et al., 2014; Hahn et al., 2006; Lee et al., 2016; Luo et al., 2014; Walther and Strik, 2012; Weickert et al., 2012).

### Expression of NRG1 type III is uncoupled from major myelin gene transcription and myelin protein expression

We further assessed the molecular mechanism responsible for the increase in myelin thickness and the functional improvements we observed. We first confirmed that the active from of NRG1 type III was expressed in Nrg1t3 and Nrg1t3;Q215X adult mice in spinal cord lysates (Fig. EV3A). Despite this, the level and activation of NRG1 type III receptors in Schwann cells, tyrosine kinase receptors ErbB2 and ErbB3, were unchanged in the presence of transgenic expression of Nrg1t3 at P30 (Fig. EV3B). Next, we quantified the level of major compact (P0, PMP22, MBP) and non-compact myelin (MAG) proteins in one-month old nerves. Despite exogenous expression of the NRG1 type III active form, no significant increase in the major myelin proteins was detected (Fig. 3A and Fig. EV3C). To further confirm that the increase in myelin thickness in Nrg1t3 animals was independent from an increase in myelin gene expression, we measured P0 protein levels in the myelin sheath using immunoelectron microscopy. We observed a decrease of P0 density in the myelin sheath of Nrg1t3 mutants (Fig. 3B), corresponding to an increase in myelin area without any increase in P0 protein levels. Thus, P0 proteins are more diluted in the myelin of Nrg1t3 mutants. This quantification supports our biochemical results of unchanged level in P0 protein despite the hypermyelination (Fig. 3A and Fig. EV3C).

**Figure 3.**
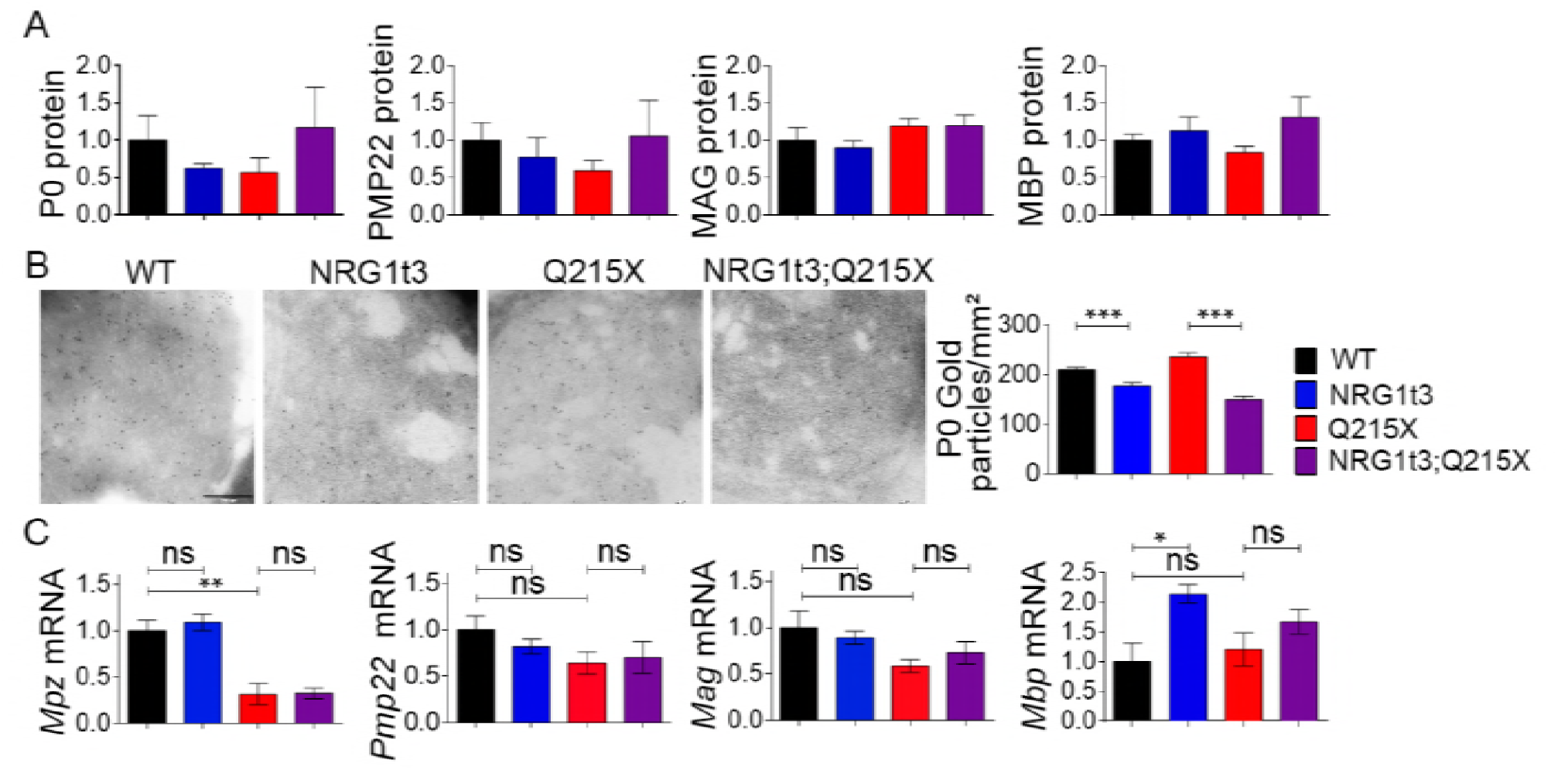
Expression of HA-NRG1 type III does not affect myelin gene expression. (**A**)Densitometries from western blot analyses of P0, PMP22, MAG and MBP performed from WT, Nrg1t3, Q215X and Nrg1t3;Q215X sciatic nerves lysates at one month of age. Protein levels remain steady in all genotypes. Representative blots are shown in Fig. EV3C. (**B**) Immunogold labeling of P0 protein from cross sections of WT, Nrg1t3, Q215X and Nrg1t3;Q215X sciatic nerves at one month. Quantification of gold particles associated with P0 from cross sections of sciatic nerve at P30. Scale bar, 200 nm (**C**) Relative quantification of mRNA *for Mpz*, *Pmp22*, *Mag* and *Mbp* in WT, NRG1t3, Q215X, NRG1t3;Q215X sciatic nerves at one month. Analyses were performed using three mice per genotype. Error bars indicate s.e.m. Statistical analyses were performed using one-way ANOVA with Bonferroni’s Multiple comparison test. * *P* < 0.05, ** *P* < 0.01, *** *P* < 0.001.

EGR2 is the key regulator of myelination, myelin protein and cholesterol/lipid biosynthesis gene expression (Le et al., 2005; Leblanc et al., 2005; Srinivasan et al., 2012; Topilko et al., 1994). Consistent with unchanged myelin gene transcription in hypermyelinated sciatic nerves of Nrg1t3 mice, *Egr2* expression was not altered in Nrg1t3 mice at one month of age (Fig EV3D-E). We also crossed our mice with mice expressing the luciferase reporter gene mRNA (*Lucif*) under the control of the Hsp68 promoter fused to a PMP22 enhancer with EGR2/SOX10 binding sites (Luc D hemizygotes (Luc D) mice, (Jones et al., 2011)). Quantification of the relative expression of Luc D in Q215X;LucD; Nrg1t3;Luc D and Nrg1t3;Q215X;LucD sciatic nerves at P15 indicated that EGR2 transcriptional activity was not affected (Fig. EV3F). These results suggest that exogenous HA-NRG1 type III promotes an increase in myelin thickness independently of the most common myelin genes and the modulation of the master *Egr2* transcription factor.

### The signaling pathways downstream of NRG1 type III are activated in Nrg1t3 mutant mice

We next sought to determine if increased NRG1 type III signaling in Nrg1t3 mice elicits activation of known signaling pathways downstream of ErbB2/3 that are necessary for Schwann cell development, differentiation and myelination, PI3K/AKT, calcineurin/NFATc4 and MAPK/ERK (Grossmann et al., 2009; Kao et al., 2009; Maurel and Salzer, 2000; Newbern et al., 2011; Ogata et al., 2004). These three signaling pathways are thought to trigger *Egr2* dependent gene activation and subsequently myelination downstream of NRG1 type III.

The activity of AKT was increased in both Nrg1t3 and Nrg1t3;Q215X one-month old sciatic nerves (Fig. 4A) which correlates with previous *in vitro* and *in vivo* observation (La Marca et al., 2011; Syed et al., 2010; Taveggia et al., 2005). In addition, we found hyperactivation of ERK1/2 pathway in both Nrg1t3 mutant mice (Fig. 4B), in agreement with recent findings that sustained and augmented MAPK/ERK signaling increases myelination (Ishii et al., 2013; Sheean et al., 2014).

The total amount and the active form of NFATc4 were unchanged (Fig. 4C). Altogether these results suggest that NRG1 type III overexpression induces the activation of both PI3K/AKT and MAPK/ERK, independently of the transcription factor EGR2.

**Figure 4.**
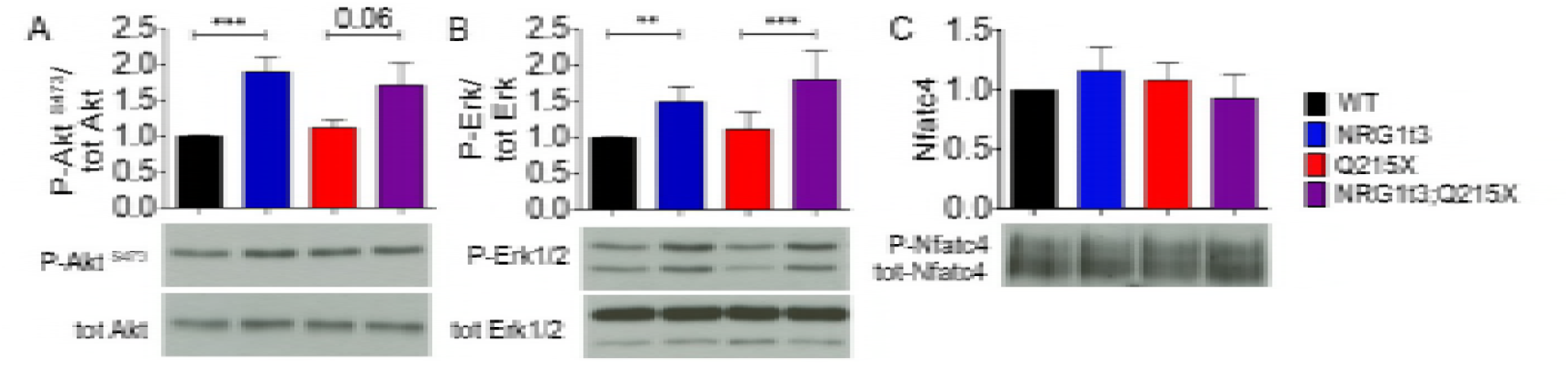
Expression of HA-NRG1 type III triggers the activation of ERK and AKT. Densitometries and western bot analyses of (**A**) P-AKT, AKT, (**B**) P-ERK, ERK, (**C**) P-NFATc4 and NFATc4 were performed on WT, Nrg1t3, Q215X and Nrg1t3;Q215X sciatic nerves lysates at one month. Increased activation of AKT and ERK was observed in NRG1t3 and Nrg1t3;Q215X. NFATc4 activation level was unchanged. Western blots were cropped. Analyses were performed using three mice per genotype. Error bars indicate s.e.m. Statistical analyses were performed using one-way ANOVA with Bonferroni’s Multiple comparison test. * *P* < 0.05, ** *P* < 0.01, *** *P* < 0.001.

### MAPK/ERK is enhanced by the exogenous expression of axonal NRG1 type III

While myelin thickness was increased with NRG1 type III transgene expression, EGR2 was unaltered. Therefore, we asked whether we could identify the transcriptomic signature of one of the canonical pathways downstream of NRG1 type III that could explain the increase in myelination in sciatic nerves of Nrg1t3 and Nrg1t3;Q215X mice.

We conducted a transcriptomic analysis of wildtype, Q215X, Nrg1t3 and Nrg1t3;Q215X sciatic nerves at 28 days of age, using mRNA next-generation sequencing. We looked for available gene libraries of signaling pathways downstream ErbB2/3/NRG1 type III, such as EGR2, NFATc4 and MAPK/ERK. We then systematically calculated the Pearson’s correlation coefficient (r) as measure of linear correlation on fold change expression between our subset of gene regulated downstream of NRG1 type III and subset of gene known to be regulated downstream of NRG1 type III-related pathways.

We first used a list of gene regulated downstream of an hypomorphic *Egr2* allele (*Egr2*^Lo/Lo^) expressed in mouse sciatic nerve at one month old (Le et al., 2005), that mimics a Egr2 loss of function. Thus, a negative correlation between Nrg1t3 and *Egr2*^Lo/Lo^ mice transcriptome should reveal the activation of EGR2 downstream of ErbB2/3/NGR1 type III signaling pathway. The absence of correlation, indicated by a coefficient r = 0.27 between *Erg2*^Lo/Lo^ and Nrg1t3 and r = 0.19 between *Erg2*^Lo/Lo^ and Nrg1t3;Q215X (Fig. 5A), confirmed that the hypermyelination mediated by HA-NRG1 type III was uncoupled from EGR2-regulated genes.

**Figure 5.**
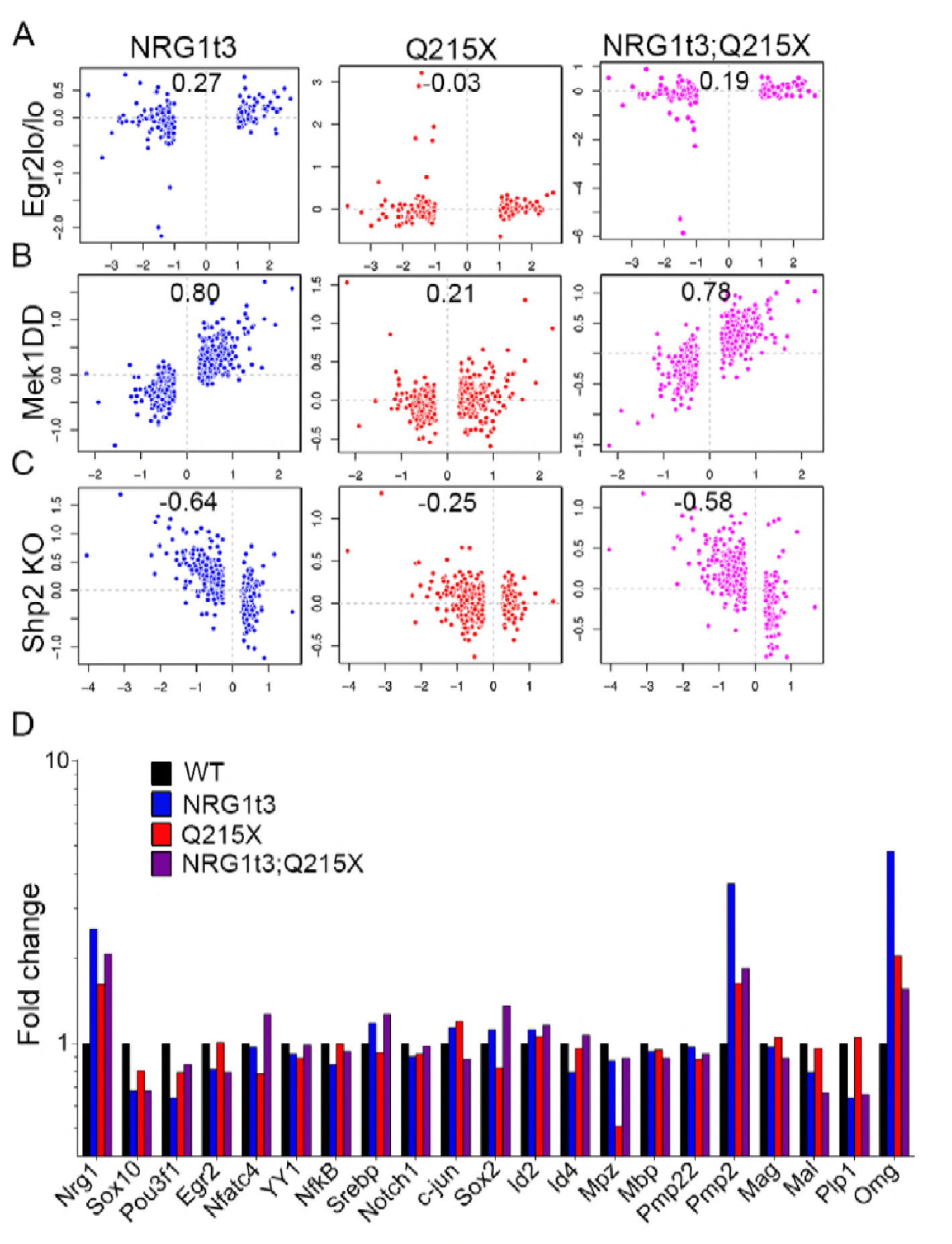
Transcriptomic signature in Nrg1t3 sciatic nerves correlates with the signature of activation of the MAPK/ERK pathway. (**A-C**) Pearson’s correlation between RNA deep sequencing from one-month old sciatic nerve of NRG1t3 (left panel in blue), Q215X (middle panel in red) and Nrg1t3;Q215X (right panel in purple) and targets downstream of EGR2 and ERK. (**A**) Pearson’s correlation analysis with combined ChIP-Sequencing and arrays data sets obtained from mice homozygous for an *Egr2* hypomorphic allele (Egr2^lo/l^°). Pair-wise correlation analysis showed no correlation between the exogenous expression of Nrg1t3 and EGR2 downstream regulated genes. (**B**) Pearson’s correlation analysis with arrays data sets obtained from conditional Schwann-specific constitutive active form of MEK1 (Mek1DD;Egr2Cre), that phosphorylates ERK. Pair-wise correlation showed a high correlation between the exogenous expression of NRG1 type III and MEK1 downstream regulated genes (Panel B blue and purple). (**C**) Pearson’s correlation analysis with arrays data set generated from Schwann cell conditional knockout for *Shp2* (Egr2Cre), which dephosphorylates ERK. Paired-wise correlation showed an anticorrelation between exogenous expression of HA-NRG1 type III and *Shp2* loss of function (panel C blue and purple). (**D**) Fold change of transcripts for neuregulin 1,transcription factors regulating myelination and myelin genes obtained from mRNA deep sequencing in Nrg1t3, Q215X and Nrg1t3;Q215X in one-month mice sciatic nerve. The quantification showed an overall unchanged level of the major regulators and components of the myelin, apart for *Pmp2* (a fatty acid binding protein, *Fabp8*) and *Omg* (oligodendrocyte myelin glycoprotein).

Second, although NFATc4 is thought to regulate *Sox10* and *Egr2* expression (Kao et al., 2009), no comprehensive analysis of NFATc4 target genes in Schwann cells has been published. Given that NFATc4 is a nuclear partner of SOX10 (Kao et al., 2009) and that SOX10 is a transcriptional activator of myelination, we used a hypergeometric-based test and data mining to first assess whether the sets of regulated genes from RNA sequence analysis of experimental nerves were enriched for genes identified by ChIP-Seq analysis of enhancers bound by SOX10. Previous studies had localized peripheral nerve enhancers through ChIP-seq of histone H3K27 acetylation, and Sox10-bound enhancers were identified (Hung et al., 2015; Lopez-Anido et al., 2015; Srinivasan et al., 2012). Sox10-bound enhancers were analyzed for the presence of NFAT motifs and annotated to the nearest expressed gene in peripheral nerve. By dividing these putative SOX10 target genes into two categories—either with or without NFATc4 motifs within 50Kb up or downstream of the gene, we inferred putative NFATc4 regulation. This analysis showed that genes regulated in nerves from either Nrg1t3 versus WT, or NRg1t3:Q215X versus Q215X, were enriched specifically for SOX10 target genes with NFATc4 motifs (Appendix Table S1). Surprisingly, those genes were significantly downregulated (Appendix Table S1), indicating that exogenous expression of HA-NRG1 type III activates less, or may even repress, NFATc4/SOX100 target genes.

Finally, to assess if genes known to be regulated downstream of the MAPK/ERK pathway correlate with those regulated downstream of ErbB2/3/NRG1 type III, we used an expression profile obtained from one-month old transgenic mice expressing a Schwann cell specific constitutive active form of the kinase MEK1 (Mek1DD) which promotes the sustained activation of ERK1/2 (Ishii et al., 2014; Sheean et al., 2014). Pearson’s correlation analysis between the two gene lists showed a striking correlation, with a coefficient r = 0.8 between Mek1DD and Nrg1t3 and r = 0.78 between Mek1DD and Nrg1t3;Q215X (Fig. 5B). No correlation was found between Q215X and Mek1DD (Fig. 5B), indicating no contribution of the P0 mutant protein itself to the regulation of MEK1-ERK pathway.

We further confirmed the activation of ERK-related genes downstream of NRG1 type III, as we found a negative correlation between our subsets and the phosphatase *Shp2* KO gene subset, an upstream regulator of ERK1/2 phosphorylation (Grossmann et al., 2009) (Fig. 5C).

Our transcriptomic results showed that among the known canonical pathways downstream of ErbB2/3/NRG1 type III, the activation of MAPK/ERK is highly correlated to an increased level of NRG1 type III. Among the genes downstream the MAPK/ERK activation pathway we found the up-regulation of fatty acid basic protein Pmp2 (encoded by *Fabp8*) and of myelin oligodendrocytes glycoprotein (encoded by *Omg*) (Fig. 5D). As furthermore, we found that none of the major myelin genes such as *Mpz, Pmp22, Mag and Mbp* as well as *Egr2* were found deregulated in Nrg1t3 transgenic mice (Fig 5D). To summarize, our analyses revealed that the gene molecular signature downstream of NRG1 type III transgenic expression is very similar to the constitutive activation of MAPK/ERK pathway. Altogether these results suggest that axonal HA-NRG1 type III expression stimulates myelin growth in an ERK dependent manner, and activates myelin genes such as *Pmp2* and *Omg* (Mikol et al., 1990; Zenker et al., 2014).

### MAPK/ERK constitutive activation in Schwann cells increases myelination in a model of congenital hypomyelination

Our data indicated that the ERK pathway is triggered downstream of the exogenous expression of HA-NRG1 type III. Therefore, we assessed whether the Schwann cell specific activation of the MAPK/ERK pathway could also restore myelin functionality in the Q215X neuropathic context. The expression of a constitutively activated form of MEK1 was restricted to Schwann cells by breeding transgenic mice hemizygous for Mek1DD (Ishii et al., 2013) with P0-Cre mice (Mek1DD;P0Cre). Morphological and ultrastructure analyses from cross sections of sciatic nerves of one-month old animals showed a strong increase in myelin thickness in Mek1DD;P0Cre;Q215X sciatic nerves, with a g-ratio of 0.39±0.002 in Mek1DD;P0-Cre (Fig. 6A, B), which is even lower than Nrg1t3;Q215X animals (Fig. 2B). Similarly to Nrg1t3 sciatic nerves, we observed that small caliber axons were predominantly hypermyelinated with Mek1DD;P0Cre, and present numerous small caliber myelinated axons (Fig. 6C and Fig. EV4). Also, no defect in myelin compaction were observed in Mek1DD;P0Cre mutants at one month (Fig. 6F-G).

Surprisingly, nerve conduction velocities were significantly reduced in Mek1DD;P0Cre mutants (Fig. 6D), and the distal amplitude of Mek1DD;P0Cre;Q215X were reduced, suggesting an axonal loss (Fig. 6E). Overall, MAPK/ERK pathway activation seems to have contrasting effects depending on its mode of activation. Beneficial effect on myelination were seen downstream of HA-NRG1 type III transgenic expression, while the direct constitutive activation of MAPK/ERK pathway showed increased myelin thickness with detrimental effect on myelin functionality.

**Figure 6.**
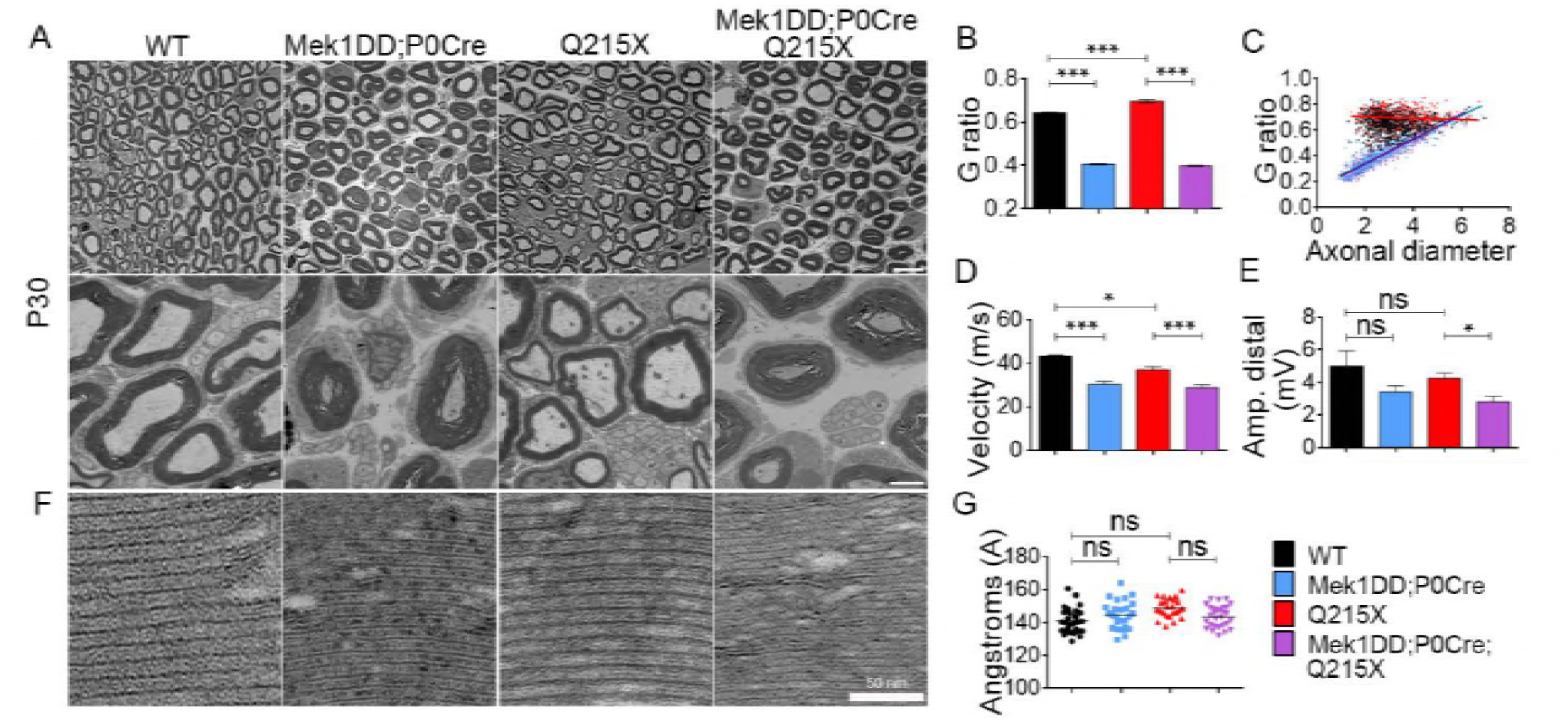
Constitutive activation of MAPK/ERK increases peripheral myelination in normal and Q215Xgenetic context, but it is functionally detrimental. (**A**) Semithin crosssections stained with toluidine blue (top panel) and electron micrographs (bottom panel) from wildtype WT, Q215X, Mek1DD hemizygote and P0-Cre hemizygote (Mek1DD;P0Cre) and Mek1DD hemizygote, P0-Cre hemizygote and Q215X (Mek1DD;P0Cre;Q215X) sciatic nerves at one month. Mek1DD;P0Cre and Mek1DD;P0Cre;Q215X fibers are hypermyelinated and Q215X are hypomyelinated. Scale bars, 10 μm (top panel) and 2 μm (bottom panel). (**B**) g-ratio was increased in Q215X (0.70±0.008) and decreased in Mek1DD;P0Cre (0.40±0.002) and Mek1DD;P0Cre;Q215X (0.39±0.002) compared to WT (0.64±0.002), indicating hypomyelination and strong hypermyelination, respectively. (**C**) g-ratio as a function of axonal diameter showed preferential hypermyelination of small fibers in Mek1DD;P0Cre and Mek1DD;P0Cre;Q215X. (**D-E**) Neurophysiological studies showed significantly reduced nerve conduction velocity in all mutants (**D**). Note that Mek1DD;P0Cre animals displayed an even more dramatic reduction. (**E**) The amplitude of compound muscle action potentials was decreased in Mek1DD;P0Cre;Q215X, suggesting axonal loss. (**F-G**) Myelin compaction (**F**) and periodicity (**G**) were not altered in Mek1DD;P0Cre and Mek1DD;P0Cre;Q215X sciatic nerves. Scale bar, 50 nm. Analyses were performed using three mice per genotype. Error bars indicate s.e.m. Statistical analyses were performed using one-way ANOVA with Bonferroni’s multiple comparison test. * *P* < 0.05, ** *P* < 0.01, *** *P* < 0.001.

### *HA-Nrg1 type III* expression promotes changes in myelin composition

Similarly to the constitutive activation of MEK1 (Mek1DD;P0Cre or Mek1DD;Egr2Cre) (Sheean et al., 2014), the transgenic expression of HA-NRG1 type III increased myelin thickness independent of the major myelin gene transcription factor EGR2 and of myelin protein synthesis (Fig. 5D and Fig. 7A). However, among the genes correlated to MAPK/ERK activation pathway, we found increased levels of *Pmp2* transcript and protein (Fig. 7A, Fig. EV5A, B), a myelin protein with functions in lipid homeostasis in myelinated Schwann cells (Zenker et al., 2014). As in several previous studies (Gillespie et al., 1990; Kadlubowski et al., 1984; Trapp et al., 1979; Zenker et al., 2014), we observed that cellular expression of PMP2 in wild type cross sections of sciatic nerve was found in compact myelin but also had cytoplasmic expression with a mosaic distribution between fibers (Fig. EV5B). In the presence of HA-NRG1 type III transgenic expression, PMP2 level was clearly increased while maintaining its mosaic distribution in sciatic nerve cross section at one month (Fig. EV5B). The specific increase in PMP2 suggests that NRG1 type III overexpression may determine a change in myelin protein composition.

**Figure 7.**
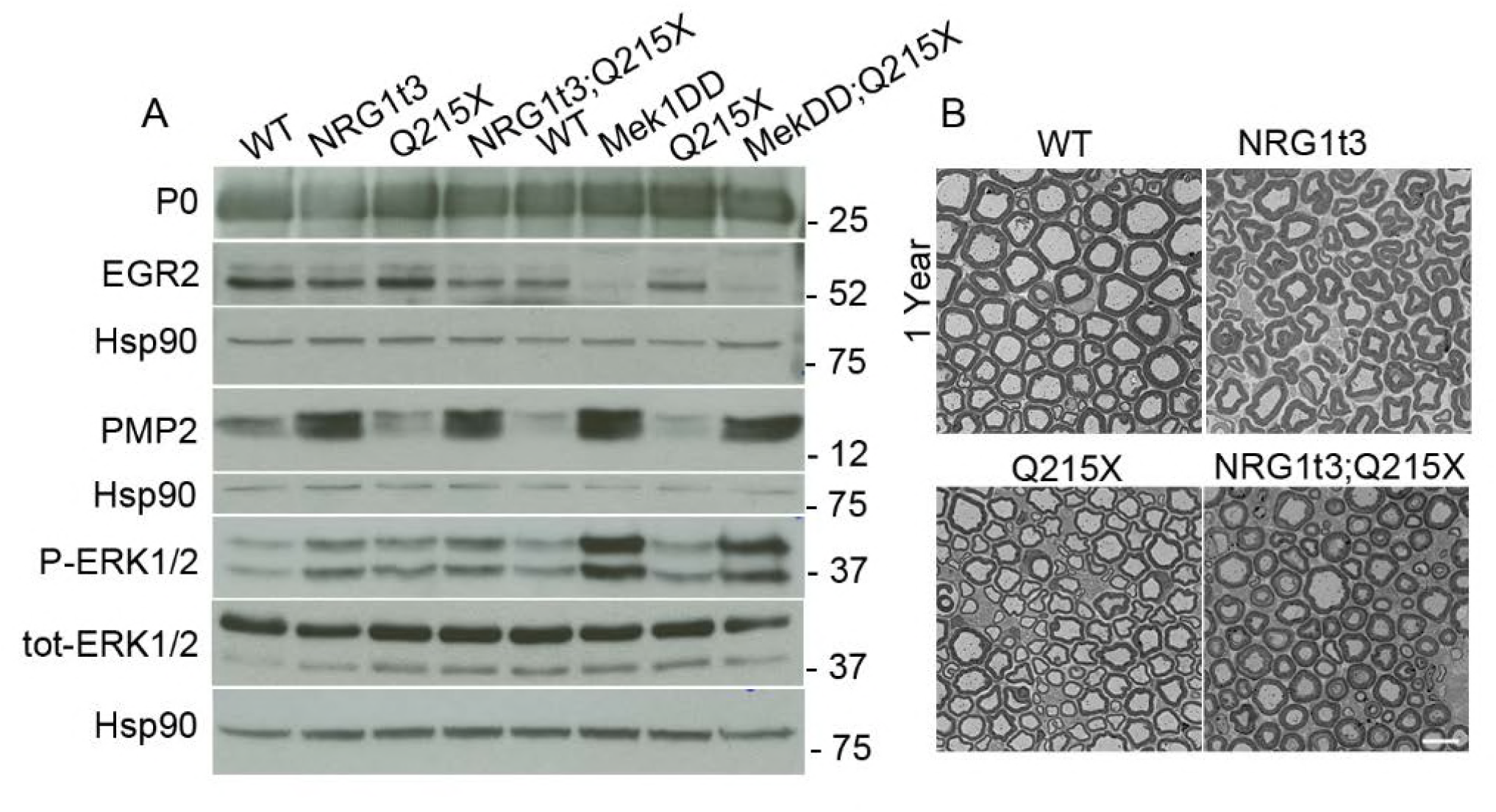
Expression of HA-NRG1 type III transgene increases PMP2 protein level. (**A**) Western blot analysis from WT, Nrg1t3, Q215X, Nrg1t3;Q215X, Mek1DD;P0Cre and Mek1DD;P0Cre;Q215X sciatic nerves at one month. Expression of HA-NRG1t3 transgene in sciatic nerves or constitutive activation the MAPK/ERK pathway in Schwann cells leads to an increase in ERK phosphorylation, a decrease in EGR2 protein levels and an increase in PMP2 protein levels. These effects are enhanced in Mek1DD;P0Cre mutants as compared to HA-NRG1t3 mutants. Western blots were cropped. (**B**) Semithin cross-sections stained with toluidine blue (top panel) and electron micrographs (bottom panel) from wildtype (WT), Nrg1t3 hemizygote (Nrg1t3), P0Q215X heterozygote (Q215X) and Nrg1t3 hemizygote and P0Q215X heterozygote (Nrg1t3;Q215X) sciatic nerves at one year.

In conclusion, both elevated NRG1 type III expression and MEK1 activation increase ERK phosphorylation but have different outcomes on myelin structure. The activation of MAPK/ERK pathway results in hypermyelination associated to nerve conduction velocity deficits (Fig. 6D) and its prolonged activation results in the formation of aberrant myelin structure i.e misfolded myelin structures increasing with age, onion bulbs, compressed axons (Ishii et al., 2016; Sheean et al., 2014). On the contrary, sustained HA-NRG1 type III transgenic expression for one year did not alter nerve physiology, and we did not observe any abnormal myelin structure (Fig. 2D, Fig. 7B), as previously reported (Michailov et al., 2004; Velanac et al., 2012).

## Discussion

In this study, we explored a suitable treatment for a severe early-onset Charcot Marie Tooth disease subtype, Congenital Hypomyelinating Neuropathy. We developed and characterized a novel mouse model of CHN, associated with a truncation of P0 cytoplasmic tail (P0Q215X) (Fratta et al., 2018). We showed that animals harboring the Q215X mutation present decreased expression of major myelin genes and proteins and displayed the hypomyelinating features of a human CHN.

In the peripheral nervous system, extrinsic signals from axonal neuregulin 1 type III regulate Schwann cell fate and myelination (Taveggia et al., 2005). Thus, the modulation of neuregulin 1 signaling is considered as a possible approach for the restoration of myelination defects in myelin diseases (Bolino et al., 2016; Fledrich et al., 2014). NRG1 type III promotes myelination by stimulating several signaling pathways, including PI3K/AKT, calcineurin/NFATc4 and MAPK/ERK, which are thought to converge on the activation of EGR2 (Fig. 8). These pathways are tightly regulated. PI3K/AKT or MAPK/ERK pathways are both necessary for myelin protein expression (Domenech-Estevez et al., 2016; He et al., 2010; Napoli et al., 2012) and imbalances in PI3K/AKT and MAPK/ERK pathways have been reported in two different demyelinating neuropathies, Charcot-Marie-Tooth type 1A (Fledrich et al., 2014; Martini, 2014) and type 1X (Groh et al., 2015). Conversely, sustained activation of either PI3K/AKT or MAPK/ERK pathways leads to hypermyelination in part through the enhancement of protein synthesis (Domenech-Estevez et al., 2016; Ishii et al., 2016; Sheean et al., 2014).

**Figure 8.**
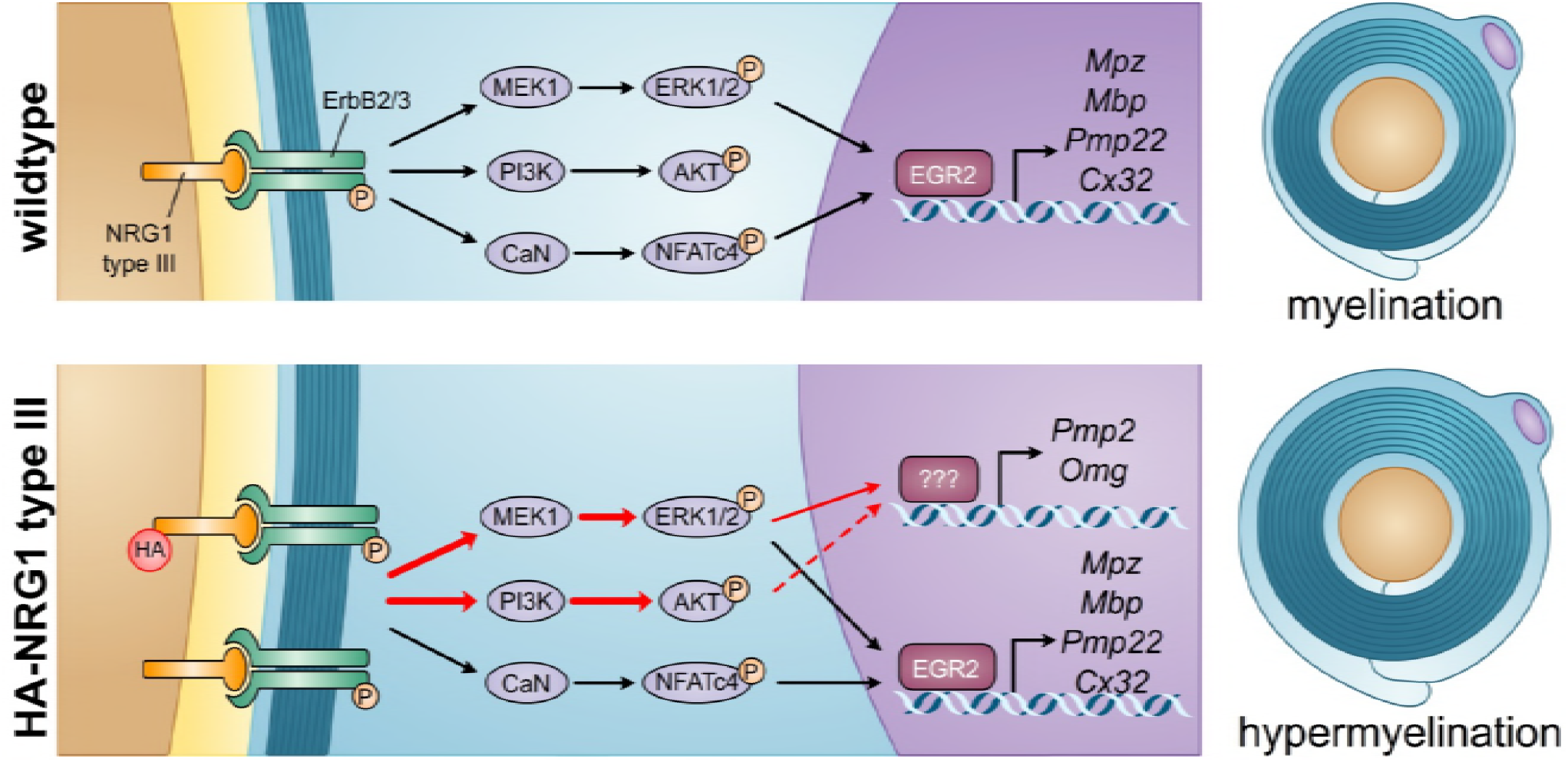
Regulation of myelination by NRG1 type III. In wildtype animals, NRG1 type III/ ErbB2/3 promotes myelination by stimulating several signaling pathways, including PI3K/AKT, MAPK/ERK and calcineurin (CaN)/NFATc4, which are thought to converge on the activation of EGR2. In Nrg1t3 animals the exogenous expression of NRG1 type III leads to the activation of both PI3K/AKT and MAPK/ERK pathways and the increased transcription of *Pmp2* and *Omg*.

Here, we show that in two different mouse models of hypomyelinated diseases i.e. congenital hypomyelinating neuropathy (this paper) and CMT1B (see accompanying paper by Scapin et al.), no initial alteration in PI3K/AKT and MAPK/ERK levels or activation was observed. Therefore, we expected that expression of exogenous NRG1 type III could improve functional outcomes in these pathologies. Indeed, we obtained the first evidences that in a CHN animal model, overexpression of NRG1 type III leads to increased activation of both PI3K/AKT and MAPK/ERK pathways, triggered the expression of genes restricted to the MAPK/ERK pathway and increases myelin thickness. We then showed that the sustained activation of MEK1-ERK in Schwann cell promotes an even stronger increase in myelin thickness, but accompanied by an alteration in nerve conduction velocity in both normal (as suggested in previous studies (Ishii et al., 2016; Sheean et al., 2014) and pathologic contexts, providing further evidences for the contrasting effect of ERK phosphorylation depending on its level of expression (Ishii et al., 2016). Finally, while NRG1 type III overexpression and MEK1-ERK sustained activation increased myelin thickness, both were uncoupled from increased expression of EGR2 or major myelin genes and proteins.

Through a comprehensive transcriptomic analysis of signaling downstream of neuregulin 1 type III in the context of neuropathy, we documented that in NRG1 type III overexpressing nerves, genes regulated by the MAPK/ERK pathway were upregulated and may contribute to increased myelin thickness in Schwann cells. Among these genes we identified, *Pmp2*, encoding for the peripheral myelin protein 2 and *Omg*, encoding for the oligodendrocyte myelin glycoprotein (OMG). *Pmp2* is expressed in myelinating cells (Patzig et al., 2011), but only some myelinated fibers appear to maintain PMP2 (Trapp et al., 1979; Zenker et al., 2014). PMP2 seems to have a role in lipid homeostasis of myelinating Schwann cells, with recent evidences that PMP2 binds fatty acids (Majava et al., 2010; Zenker et al., 2014). This is particularly interesting as NRG1 was also identified upstream of MAF, a regulator of cholesterol biosynthesis (Kim et al., 2018), and in mice overexpressing NRG1 type III, cholesterol along with saturated and unsaturated fatty acids are increased (see accompanying paper Scapin et al.). On the other hand, there are conflicting reports on the expression of OMG in Schwann cells (Apostolski et al., 1994; Chang et al., 2010; Huang et al., 2005), and the function of OMG is unknown. Total knockout mice revealed that both PMP2 and OMG are dispensable to the formation or maintenance of myelin structure in the peripheral and central nervous system, respectively (Chang et al., 2010; Zenker et al., 2014). However, two recent studies in human revealed that de novo dominant mutations in *PMP2* were identified in families with type 1 Charcot Marie Tooth demyelinated disease (Gonzaga-Jauregui et al., 2015; Motley et al., 2016). Thus, it has been suggested that mutations in *PMP2* result to a probable toxic gain of function in myelinating Schwann cells causing demyelination (Gonzaga-Jauregui et al., 2015; Motley et al., 2016). Here, the hyperactivation of the MAPK/ERK pathway in Nrg1t3 animals appears to stimulate specifically the transcription of these less studied myelin proteins. The understanding of PMP2 and OMG function in Nrg1t3 animals could help us to understand further the mechanisms that drive hypermyelination.

MAPK/ERK activation has been shown to have beneficial promyelinating function during normal development (Grossmann et al., 2009; Newbern et al., 2011; Shin et al., 2014) and to promote remyelination after injury (Napoli et al., 2012). Paradoxically, the activation of MAPK/ERK in Schwann cells has been also associated to detrimental effects on Schwann cell survival, development, myelination, remyelination (Maurel and Salzer, 2000; Napoli et al., 2012; Ogata et al., 2004; Syed et al., 2010) and axonal regeneration (Cervellini et al., 2018). The discrepancies between these studies could be explained in part by the context, the timing and the level of MAPK/ERK activation. A limited activation of MAPK/ERK, as in Nrg1t3 mice, was not associated to myelin structural or functional alterations, in accordance with previous histological examination in Nrg1t3 sciatic nerve (Michailov et al., 2004; Velanac et al., 2012). However, we and others showed that a sustained activation of MEK1 is associated to aberrant myelin structures and affects axon and myelin integrity (Ishii et al., 2016; Sheean et al., 2014). Interestingly, we noticed that a strong decrease of EGR2 protein level specifically correlated with the sustained activation of MEK1 in Schwann cells, suggesting a dose effect of ERK phosphorylation levels on the regulation of EGR2 expression.

It is possible that in Nrg1t3 nerves other molecular mechanisms downstream of NRG1 type III limit MAPK/ERK activation over time, in contrast to the constitutively activated MAPK/ERK pathway. For example, we found a decreased activation of genes with NFATc4 motifs, while the total amount and the active (nuclear) form of NFATc4 were unchanged. In addition, we saw an upregulation of the PI3K/AKT pathway, which might be important to balance MAPK/ERK signaling, as suggested previously (Fledrich et al., 2014). Also, possible upstream mechanisms such as laminin and laminin receptors can limit NRG1 type III effect on myelination (Ghidinelli et al., 2017; Heller et al., 2014; Mogha et al., 2016; Monk et al., 2015).

Independently of the mechanism, we show that exogenous expression of HA-NRG1 type III has no detrimental effects on peripheral myelination and improved myelin thickness and nerve conduction velocity in both CHN and CMT1B (see accompanying paper Scapin et al.). Thus, our observations suggest that controlled activation of NRG1 type III could represent a favorable therapeutic approach in several human peripheral neuropathies associated with hypomyelination. During Schwann cell myelination, NRG1 type III activity is tightly regulated by complex and stepwise proteolytic cleavages by BACE1, TACE, GPR44 and I-CLips (Fleck et al., 2016; Hu et al., 2006; La Marca et al., 2011; Trimarco et al., 2014; Willem et al., 2006). It is therefore possible to regulate the activity of specific proteases to favor the cleavage products of NRG1 type III beneficial for myelination. By this principle, inhibition of TACE improved myelination in an hypermyelinating model CMT4B showing that modulation of NRG1 type III signalling in both directions can be beneficial in different CMT neuropathies (Bolino et al., 2016). However, the modulation of a single protease *in vivo* can be challenging because inhibition of these proteases in regions other than peripheral nerves could have unwanted corollary effects; for example, TACE inhibition might promote amyloid production in the brain, and sustained activation of MAPK/ERK in non-myelinating Schwann cells promotes corneal neurofibromas (Bargagna-Mohan et al., 2017). Finally, NRG1 type III is expressed in neurons of the central nervous system and exogenous expression of HA-NRG1 type III induces hypermyelination in the corpus callosum (Brinkmann et al., 2008; Taveggia et al., 2008). The consequences of this increase in myelination in the brain have not been fully investigated, but several studies have linked the increase of NRG1 type III to schizophrenia, possibly by disrupting cortical neuronal synaptic plasticity, and altering synaptic transmission (Agarwal et al., 2014; Hahn et al., 2006; Weickert et al., 2012). Thus, a better understanding of the function of NRG1 type III and its proteases in adult Schwann cells is required to manipulate NRG1 signaling specifically in the peripheral nervous system.

In conclusion, this study shows encouraging results for the application of NRG1 type III modulations in improving myelination in the context of peripheral hypomyelinated neuropathies.

## Materials and methods

### Animal model

All animal experimentation was performed in strict accord with Institutional Animal Care and Use Committee approved protocols of San Raffaele Institute and University at Buffalo (n. 363 and UB1188M respectively). P0Q215X knock-in mice (hereafter called Q215X) contain a targeted mutation in *Mpz* exon 5 (Fratta et al., 2018). Neuregulin 1 type III hemagglutinin-tagged transgenic mice (hereafter called Nrg1t3), expressing *Nrg1 type III* under the control of neuronal promoter Thy1.2, were previously described (Velanac et al., 2012). The R26Stop^FL^Mek1DD mice (hereafter called Mek1DD), expressing conditionally an activated form of MEK1, were previously described (Srinivasan et al., 2009). Mek1DD mice were cross-bred with P0-Cre transgenic mice that express Cre recombinase starting at E13.5 (Feltri et al., 1999; Feltri et al., 2002). We used a reporter mouse model containing a basal Hsp68 promoter upstream of the luciferase reporter gene activated by the +11 kb human *PMP22* intronic enhancer containing a combinatorial Egr2/Sox10 binding site, transgenic luciferase line D (hereafter called Luc D) (Jones et al., 2011).

Q215X and P0-Cre mice were in FVB/N congenic background. Nrg1t3 and Mek1DD mice were in C57BL/6 background. F1 mice were respectively generated by crossbreeding Q215X mice with the transgenic Nrg1t3 mice and Q215X with Mek1DD in C57BL/6xFVB/N mixed background. Luciferase mice were in mixed background C57BL/6xFVB/N/CBA. Genotyping of mutant P0-Cre, Nrg1t3 and Mek1DD mice was performed by PCR on tail genomic DNA, as described previously (Feltri et al., 1999; Jones et al., 2011; Michailov et al., 2004; Srinivasan et al., 2009). The genotype of Q215X animals was determined by PCR analysis with *Mpz* forward primer: 5'-CAACCTCTCTTGCCACAGTG-3' and *Mpz* reverse primer: 5'-GCTAACCGCTATTTCTTATCC-3' which amplified 400 and 280 nucleotide fragments, for Q215X and wild type allele respectively. PCR conditions were: 94 °C for 1 minute (min), 56 °C for 1 min and 72 °C for 1 min (29 cycles), followed by 10 min extension at 72 °C, in a standard PCR reaction mix.

No animals were excluded from the study. Animals were housed in cages of no more than 5 animals in 12 h light / dark cycles. Mutant and control littermates from either sex were sacrificed at the indicated ages, and sciatic nerves were dissected.

### Functional analyses

Grid walking test was used to measure the frequency of step errors on rungs of an elevated grid. Littermate mice were allowed to walk on a mesh grid for 5 trials of 5 min each, with the first three trials considered as training period. Hindpaw placement errors (Foot faults) defined as passage of the hindpaw through an opening in the grid were determined as a proportion of total hindpaw steps on horizontal grid mesh (1.2 × 1.2 cm openings). Sciatic nerve motor conduction velocity and amplitude were obtained as previously described (Poitelon et al., 2015; Poitelon et al., 2016).

### Morphology

Semithin section and electron microscopic analyses of sciatic nerves were performed as previously described (Quattrini et al., 1996). For g-ratio (axon diameter/fiber diameter) and axonal distribution, four semithin images per sciatic nerve were acquired with the 100X objective. Three to four animals per age per genotype were analyzed. Axon and fiber diameters were quantified using the Leica QWin software (Leica Microsystem). Data were analyzed using GraphPad Prism 6.01. Periodicity was measured at 20,000X magnification as previously described (Wrabetz et al., 2006). Immuno-electron microscopy was performed as in (Quattrini et al., 1996) but with embedding modified as described (Yin et al., 2000). Primary antibody was chicken anti-P0 (Aves). Gold-conjugated secondary antibodies were 10 nM for P0 (British Biocell International). For teased fibers, sciatic nerves were fixed in 2 % glutaraldehyde overnight at 4 °C, washed 3 times in phosphate buffer (79 mM Na2HPO4, 21 mM NaH2PO4, pH 7.4). Sciatic nerves were stained in 1 % osmium, washed 4 times in phosphate buffer and incubated at 55 °C for twelve hours (h) in 30 % glycerol followed by 12 h in 60 % glycerol and 12 h in 100 % glycerol.

### Western analysis

Sciatic nerves were dissected, frozen in liquid nitrogen, pulverized and resuspended in lysis buffer (150 mM NaCl, 25 mM HEPES, 0.3 % CHAPS pH 7.4, 1 mM Na3VO4, 1 mM NaF and 1:100 Protease Inhibitor Cocktail (Roche). Protein lysates were centrifuged at 13000 g for 20 min at 4°C. Supernatant protein concentrations were determined by BCA protein assay (Thermo Scientific) according to the manufacturer’s instructions. Equal amounts of homogenates were diluted 3:1 in 4X Laemmli (250 mM Tris-HCl pH 6.8, 8 % SDS, 8 % β-Mercaptoethanol, 40 % Glycerol, 0.02 % Bromophenol Blue), denatured 5 min at 100 °C, resolved on SDS-polyacrylamide gel, and electroblotted onto PVDF membrane. Blots were then blocked with 5 % BSA in 1X PBS, 0.05 % Tween-20 and incubated over night with the appropriate antibody. Cell Signalling anti-AKT (9272) 1/1,000, anti-p-AKT Ser473 (9271) 1/1,000, anti-p-AKT Thr308 (5106) 1/1000, anti-ERK 1/2 (#9102) 1/1,000, anti-p-ERK 1/2 p44/42 (9101) 1/1,000, anti-NFATc4 (2183) 1/1000. Aves anti-P0 (PZO) 1/5000. Covance anti-MBP (SMI94) 1/1000, anti-TUJ1 (MMS-435P) 1/5000. Sigma anti-GAPDH (G9545) 1/2000, anti-Calnexin (C4731) 1/3000. Santa Cruz anti-PMP2 (sc-374058), anti-HSP90 (sc-7947) 1/1000. Abcam anti-PMP22 (ab15506) 1/1000, anti-ErbB2 (ab2428) 1/200 and anti p-ErbB2 Y1248 (ab47755) 1/1000. Invitrogen anti-MAG (34-6200) 1/1000. Roche anti-HA (11867423001) 1/200. anti-EGR2 1/200 was gifted by Dr. Meijer, Centre for Neuroregeneration, Edinburgh. Membranes were then rinsed in 1X PBS and incubated for 1h with secondary antibodies. Blots were developed using ECL, ECL plus (GE Healthcare) or Odyssey CLx infrared imaging system (Li-Cor). Western blots were quantified using Image J software (<u>http://imagej.nih.gov/ij</u>). Each experiment was performed at least two times on nerves from at least three animals.

### Immunofluorescence and immunohistochemistry

For immunohistochemistry, sciatic nerves were fixed with 4 % PFA and embedded in OCT. Cross sections were permeabilized with cold MetOH. Sections were rinsed in 1X PBS, blocked for 1 h in 20 % fetal bovine serum, 2 % BSA, 0.1 % Triton X-100 in 1X PBS. The following primary antibodies were incubated overnight: Proteintech anti-PMP2 (12717-AP) 1/500, Aves Labs anti-P0 (PZO) 1/1000. Sections were rinsed in 1X PBS, incubated 1 h with Jackson TRITC-conjugated (711-025-152) 1/200 and with Jackson DyLight 488 1/1,000, stained with DAPI, and mounted with Vectashield (Vector Laboratories). Images were acquired with a Zeiss ApoTome.

### RNA extraction and quantitative RT-PCR

Total RNA was extracted from sciatic nerves using TRIzol^®^ (Life Technologies). Several pools of nerves of two to six nerves were used depending the animal age. For RT-QPCR analysis, first strand cDNA was synthesized from 1 μg total RNA using polythymine and random primers with Superscript III RNAse H reverse transcriptase (Invitrogen). The expression of selected mRNAs was determined by quantitative real-time PCR using 20 ng of cDNA. Samples were processed in triplicate and reactions without target cDNA were used as negative control for each reaction. PCRs were performed on 96-well plates using the Power SYBR Green PCR Master Mix (Applied Biosystems), Taqman probe assays or FastStart Universal Probe Master (Roche Diagnostic) following manufacturer’s conditions. Data were analyzed using the threshold cycle (Ct) and 2('^ΔΔ^Ct) method. *Actb, Hsp90, Gapdh* or *Pgk1* genes were used as reference for normalizing the data. The Taqman probes used are the following: *Actb* (Mm00607939-S1), *Gapdh* (Mm99999915-g1), *Pgk1* (Mm00435617), *Pmp2* (Mm03015237_m1), *Mpz* (Mm00485139-m1), *Mbp* (Mm01262035), *Mag* (Mm00487541-m1), *Egr2* (Mm00456650-m1), *Pmp22* (Mm01333393_m1).

### RNAseq analysis

Sciatic nerves at P30 were dissected, frozen in liquid nitrogen and pulverized. Total RNA was prepared from pools of two nerves with TRIzol (Roche Diagnostic), then purified with RNeasy column (Qiagen). Samples were quantified using Ribogreen Assay (Invitrogen) and the quality of samples was checked using Agilent Bioanalyzer 2100 RNA nano 6000 chip (Agilent). Illumina TruSeq RNA sample preparation kit (Illumina) was used to prepare cDNA libraries from RNA samples. Samples were poly A selected to isolate mRNA, the mRNA was cleaved into fragments, the first strand reverse transcribed to cDNA using SuperScript II reverse Transcriptase (Invitrogen) and random primers, followed by second strand cDNA synthesis using Second Strand Master Mix supplied with the kit. After end repair, the addition of a single ‘A’ base, and ligation with adapters, the products were enriched and purified with PCR to create the final cDNA library as per manufacturer’s protocol. cDNA libraries were quantified using Picogreen Assay (Invitrogen) and Library Quantification kit (Kapa Biosystems). Agilent Bioanalyzer 2100 DNA 7500 chip was used to confirm the quality and size of the cDNA libraries. The cDNA libraries were then normalized, pooled and paired-end sequenced (100 standard cycles) using the Illumina HiSeq2500 following the manufacturer’s instructions at the UB Genomics and Bioinformatics Core Facility (Buffalo, NY). Reads were aligned to the mouse reference genome (mm10) with the transcript annotation (UCSC mm10 annotation download from TopHat website) using TopHat (v2.0.9). Quantification for the gene expression and differential expression analysis were done using the Cufflinks software (v2.2.0). Data are available at http://www.ncbi.nlm.nih.gov/geo/ accession number: GSE101808.

### Experimental Design

Experiments were not randomized, but data collection and analysis were performed blindly to the conditions of the experiments. Researchers blinded to the genotype performed behavioral analyses, nerve conduction velocities and morphometric analyses. This study was not preregistered. Most experiments were conducted with 3 animals per age and per genotype. No statistical methods were used to predetermine sample sizes, but our sample sizes are similar to those generally employed in the field.

### Statistical analyses

The data obtained are presented in mean ± s.e.m. One-way ANOVA with Bonferroni’s comparison test was used for statistical analysis of the differences among multiples groups. Values of *P* < 0.05 were considered to represent a significant difference.

Each Pearson’s correlation coefficient (r) was calculated from sets of genes selected with a fold change cutoff ≥ 1.2. Correlation plots show the linear relationship between gene sets obtained from P30 sciatic nerves.

A hypergeometric-based test (Fisher exact test) interrogated for enrichment of regulated genes from RNA sequencing from nerves of various genotypes, in subsets of regulated genes selected from Sox10 ChIP-Seq (Lopez-Anido et al., 2015; Srinivasan et al., 2012) with or without NFATc4 motif enrichment. Motifs (MA0152.1 Jaspar) were called using the Homer annotate Peaks program with option -m (Heinz et al., 2010). The rationale for using the hypergeometric test was that if genes activated downstream of Nrg1 type III had any biological or functional association with the calcineurin/NFATc4 pathway, a higher number of genes would be found regulated in nerve and present in ChIP-Seq libraries than expected by chance.

## Acknowledgements

We thank Edward Hurley for semithin sections and electron microscopic preparations.

This work was funded by Telethon Grant GPP10007 (L.W., M.L.F and C.T.), NINDS grant R01NS096104 (LW), Charcot-Marie-Tooth Association grant CMT1B-003 (LW) and Peripheral Nerve Society Fellowship Grant for research in neuropathy 2012-2013 (SB).

## Author contribution

S.B., L.W. designed research, analyzed and interpreted data; S.B., F.O., G.S. performed the majority of research; J.W. performed RNAseq analysis; C.L., J.S. performed motif enrichment analysis; N.S. performed electrophysiology analysis; C.S., N.R., C.W., C.T., Y.P., M.D. provided technical assistance; A.I., R.B., M.S., K.N. contributed analytical tools; S.B., Y.P., L.W. wrote the manuscript; M.L.F., L.W. provided funding and scientific direction; J.S., R.B., M.S., P.F., Y.P., M.D., M.L.F. critically reviewed the manuscript.

## Conflict of interest

The authors declare no competing financial interests.

## The paper explained

### Problem

Hereditary demyelinating neuropathies are important neuromuscular disorders with significant social and economic impact. Among them, neuropathies due to Myelin Protein Zero (*MPZ*) mutations are relatively common and of widely varying phenotype, including Charcot Marie Tooth 1B (CMT1B) with onset in childhood or in adulthood, and Congenital Hypomyelination neuropathy (CHN) apparent at birth. There is no treatment for MPZ-neuropathies. We have produced and characterized a variety of mouse models for *MPZ*-neuropathies and showed that different mutations cause pathology by different mechanisms. Thus, even within MPZ-neuropathies, it may be difficult to identify unifying therapies that target each pathogenic mechanism. However, because hypomyelination and demyelination are common features of all these neuropathies, we turned to final-common pathways that regulate myelination.

### Results

Neuregulin 1 type III (NRG1 type III) promotes peripheral myelin formation and thickness. Therefore, we explored the hypothesis that augmenting NRG1 type III levels may ameliorate hypomyelination, demyelination and inefficient remyelinaton in a preclinical model of CHN and CMT1B (Scapin et al., this issue), despite different pathogenic mechanisms.

### Impact

Our paper establishes for the first time that overexpression of NRG1 type III rescues myelination and nerve conduction velocity in mice carrying a nonsense P0 mutation that causes CHN. These effects are accomplished without altering major myelin protein levels. Instead, genes regulated by the MAPK/ERK pathway are upregulated and may contribute to increased myelin thickness in Schwann cells. Together, this study and the one led by the D’Antonio lab (Scapin et al., this issue) provide novel results supporting that exogenous expression of NRG1 type III has no detrimental effects on peripheral myelination and can improve myelination in CHN and CMT1B. Thus, our observations suggest that controlled activation of NRG1 type III could represent a favorable therapeutic approach in several human peripheral neuropathies associated with hypomyelination.

## For More information

http://www.buffalo.edu/hunter-james-kelly-research-institute.html

## Expanded View Figure Legends

**Figure EV1.**
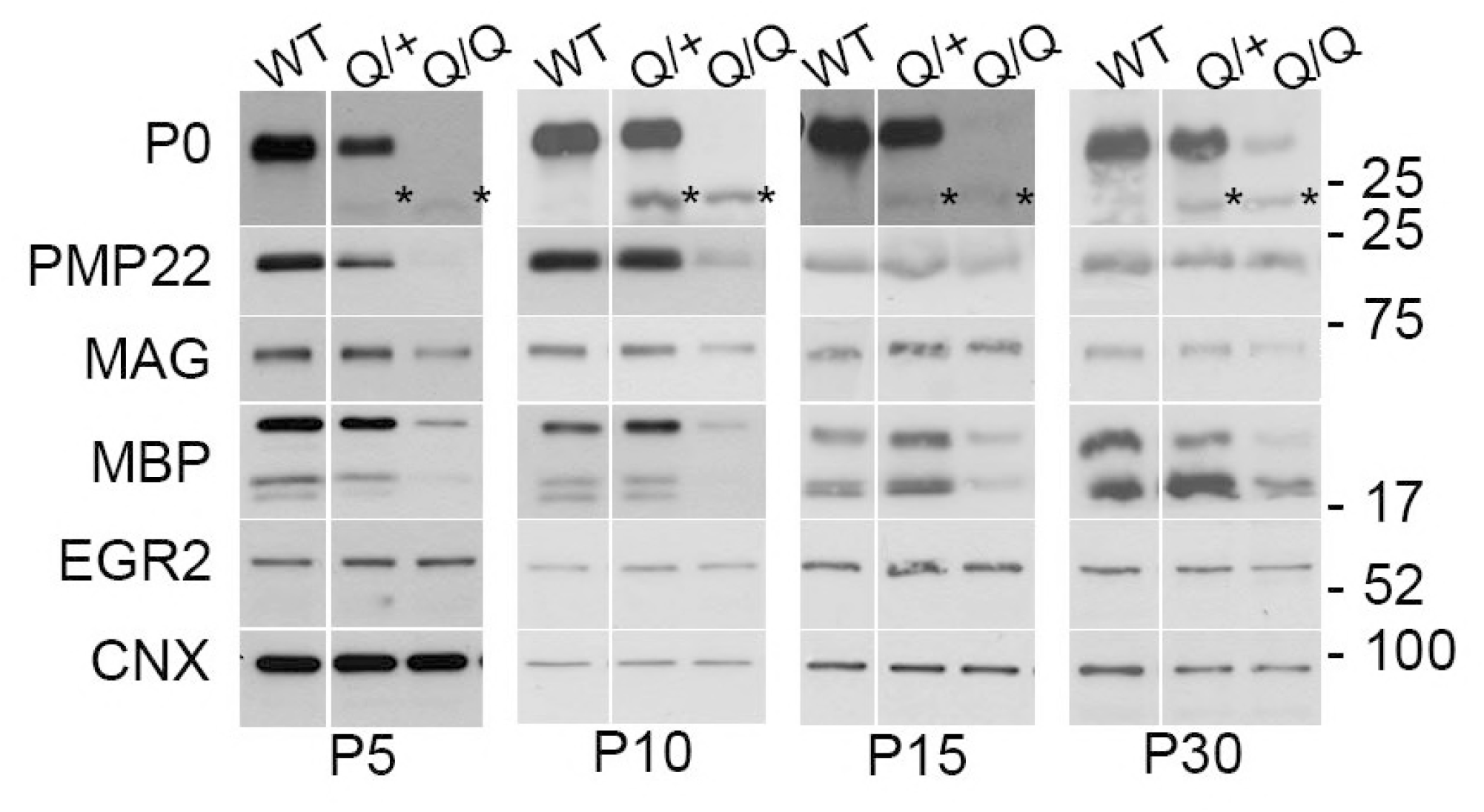
The P0Q215X disrupts myelin gene regulation and modifies myelin protein expression throughout the development. Western analysis from WT, Q215X (Q/+), Q215X/Q215X (Q/Q) sciatic nerves during development at P5, P10, P15 and P30 days. Blots were probed for P0, PMP22, MAG, MBP and EGR2. The Q215X allele cause a reduction of P0 stability (Fratta et al., 2018). Calnexin was used as loading control. Analyses were performed using three mice per genotype. Western blots were cropped.

**Figure EV2.**
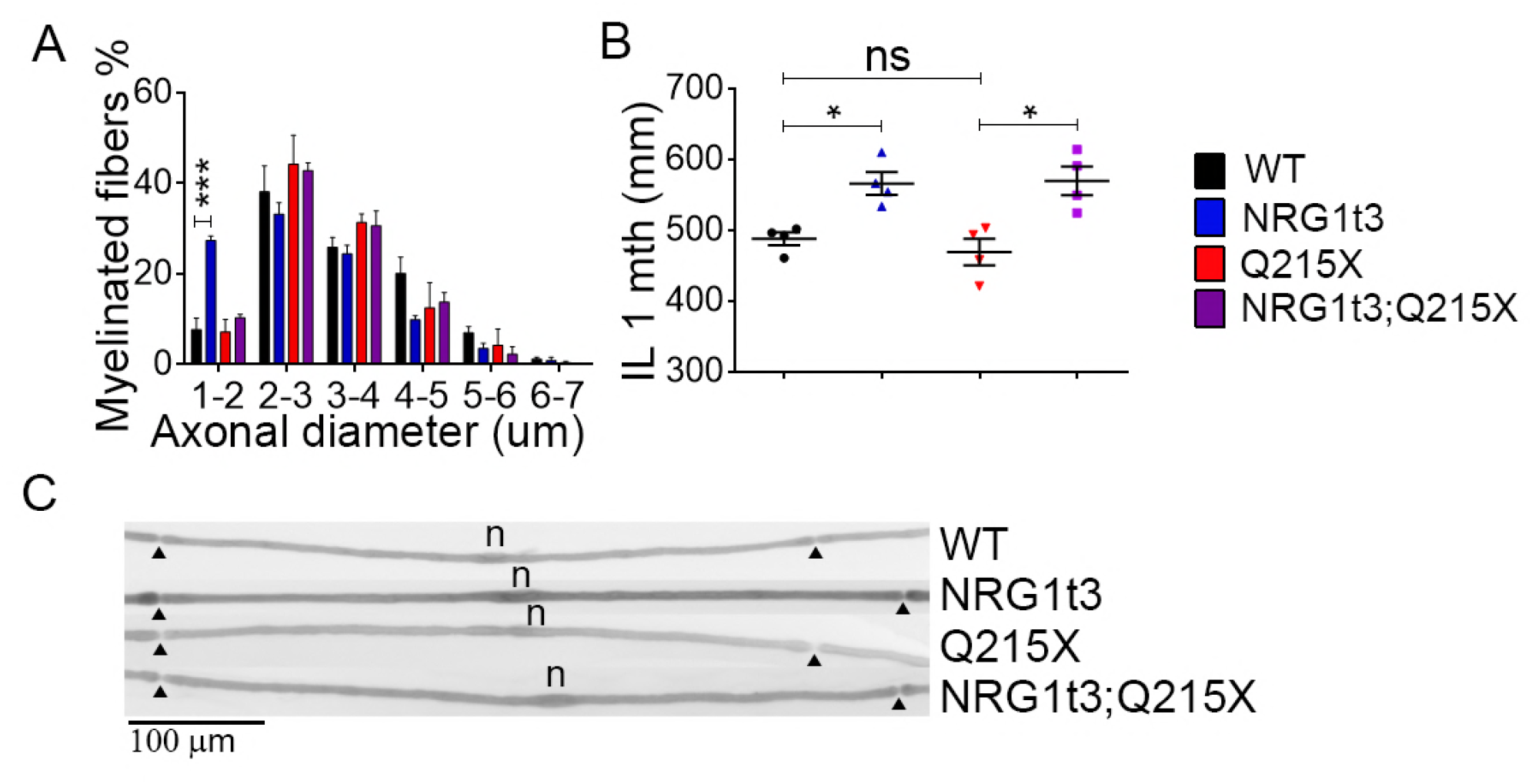
Expression of HA-NRG1 type III promotes a shift in the size of myelinated fiber population and an increase in internodal length. (A) The distribution of myelinated fibers shows a larger population of small myelinated axon (axon diameter from 1 to 2 um) in Nrg1t3 compare to WT. (**B-C**) While no reduction in internodal lengths was measured in the Q215X, the internodal lengths were increased in both Nrg1t3 and Nrg1t3;Q215X and Nrg1t3 by approximately 15% (**B**). At least 100 internodes were measured (**C**). Scale bar 100 μm. Analyses were performed using three mice per genotype. Error bars indicate s.e.m. Statistical analyses were performed using one-way ANOVA with Bonferroni’s multiple comparison test. * *P* < 0.05.

**Figure EV3.**
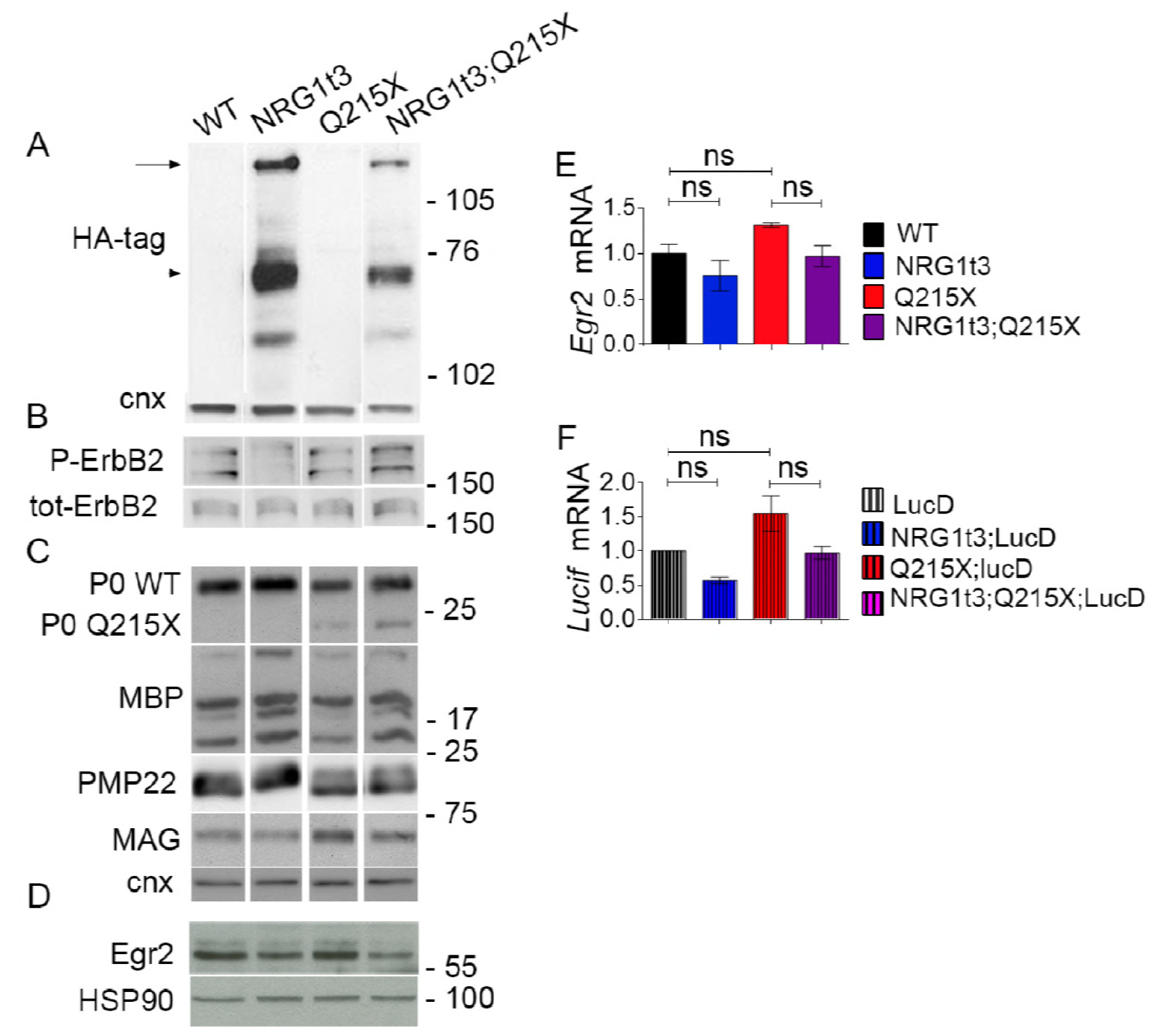
The hypermyelination mediated by HA-NRG1 type III transgenic expression does not correlates with an increase in major myelin components or in EGR2 activation. (**A**) Western blot analyses from WT, Q215X, Nrg1t3;Q215X and Nrg1t3 spinal cord at P30. Blots were probed for HA tag. HA is tagged to the NRG1 type III transgene. Calnexin was used as loading control. We detected exogenous neuregulin precursor protein level and its cleavage products. The upper band corresponds to the full-length NRG1 type III inactive isoform (135-kDa, arrow), the 75-kDa band corresponds to the active form (arrow head) of NRG1 type III produced by BACE1-cleavage, as described in the literature (La Marca et al., 2011; Velanac et al., 2012). (**B-D**) Western blot analyses from WT, Q215X, Nrg1t3;Q215X and Nrg1t3 sciatic nerves at P30. Blots were probed for P-ErbB2, ErbB2 (**B**), P0, MBP, PMP22, MAG (**C**) and EGR2 (**D**). HSP90 was used as loading control. (**E**) Relative quantification of *Egr2* mRNA from WT, Nrg1t3, Q215X, HA-Nrg1t3;Q215X sciatic nerves at P30. (**F**) Relative quantification of luciferase reporter gene mRNA (*Lucif*) from Luc D hemizygotes (Luc D), Q215X;LucD, Nrg1t3;Luc D and Nrg1t3;Q215X;LucD sciatic nerves at P15. Luc D mice harbor a combinatorial EGR2/SOX10 binding site upstream of basal *Hsp68* promoter. No significant differences were found at P15. Analyses were performed using three mice per genotype. Error bars indicate s.e.m. Statistical analyses were performed using one-way ANOVA with Bonferroni’s multiple comparison test. *** *P* < 0.001.

**Figure EV4.**
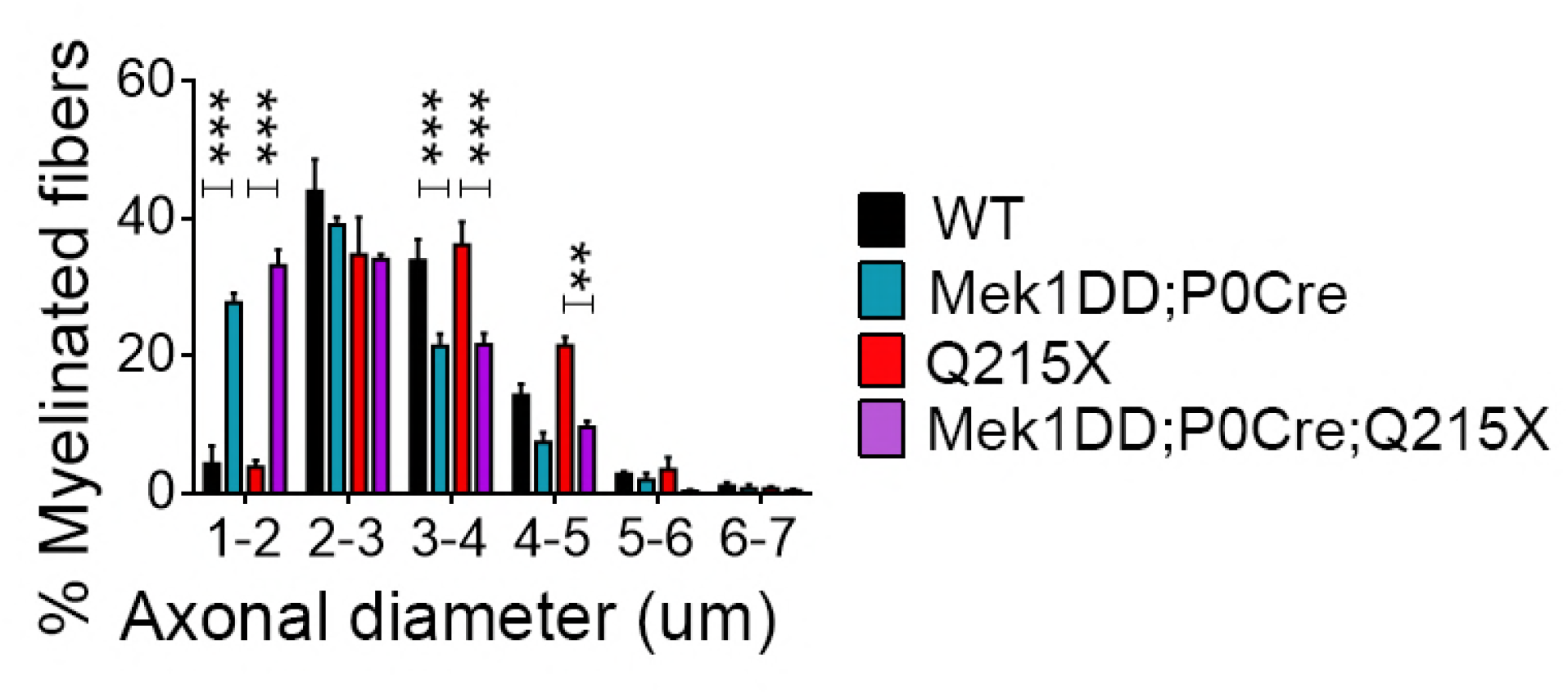
Mek1DD;P0Cre transgene promotes a shift in the size of myelinated fiber population. The distribution of myelinated fibers shows a larger population of small myelinated axon (axon diameter from 1 to 2 um) in Mek1DD;P0Cre and Mek1DD;P0Cre;Q215X as compared to WT and Q215X at one month. Analyses were performed using three mice per genotype. Western blot were cropped. Statistical analyses were performed using one-way ANOVA with Bonferroni’s multiple comparison test. * *P* < 0.01, *** *P* < 0.001.

**Figure EV5.**
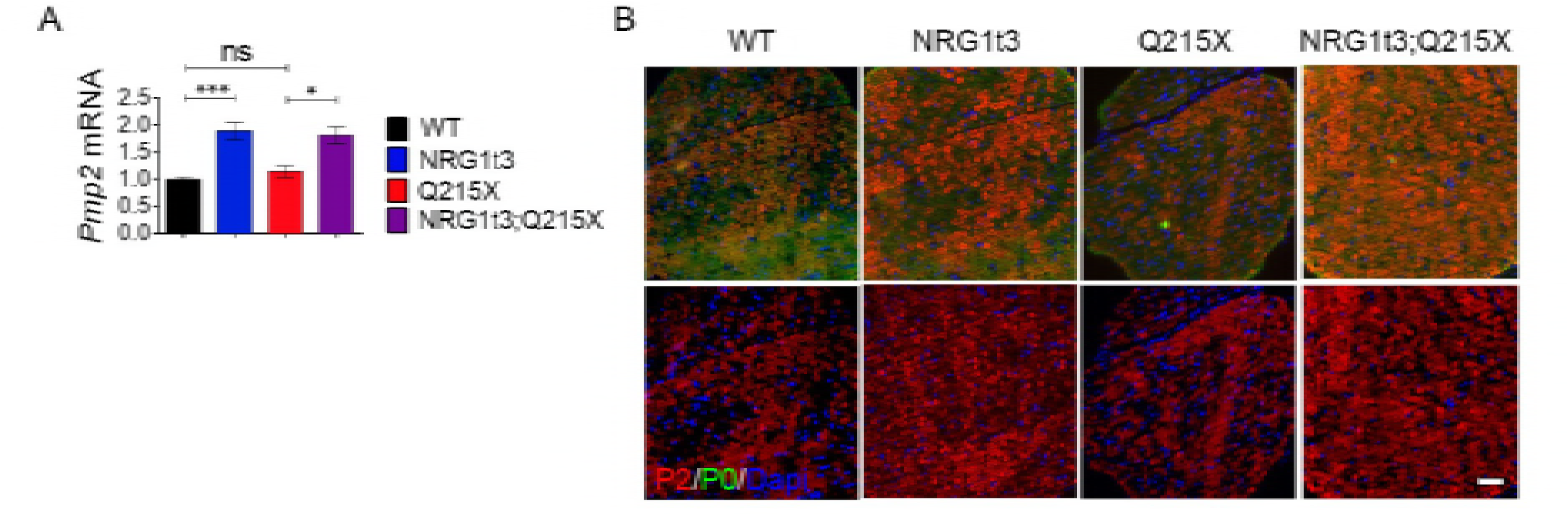
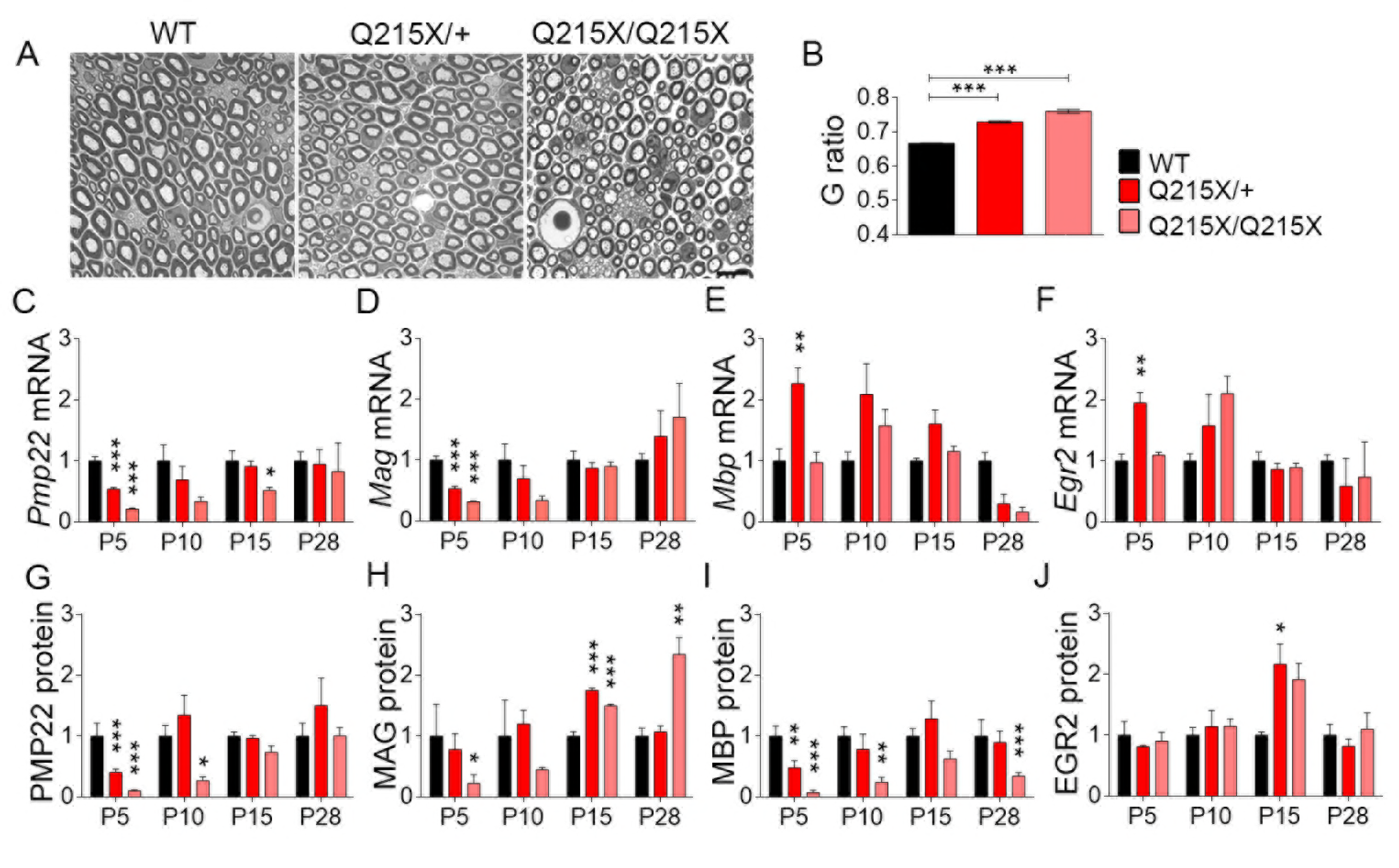
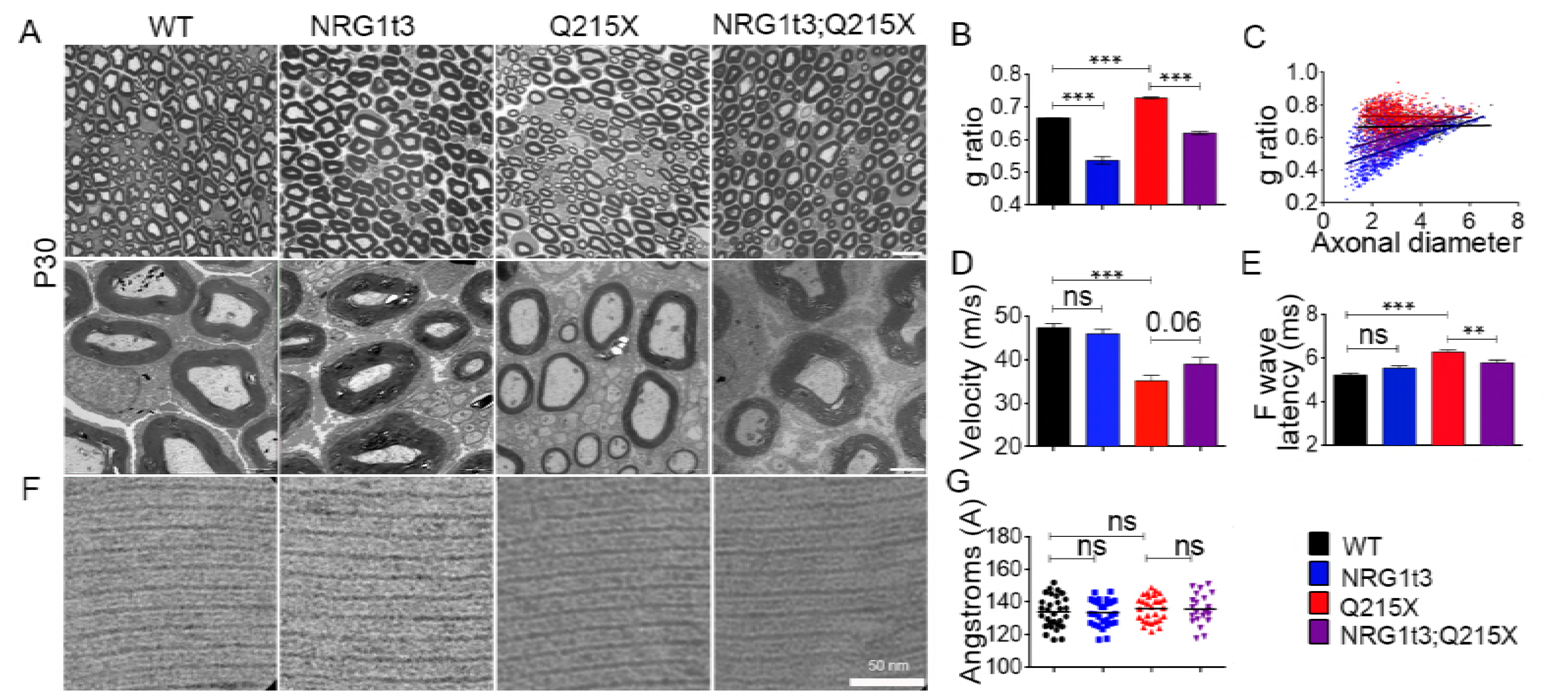
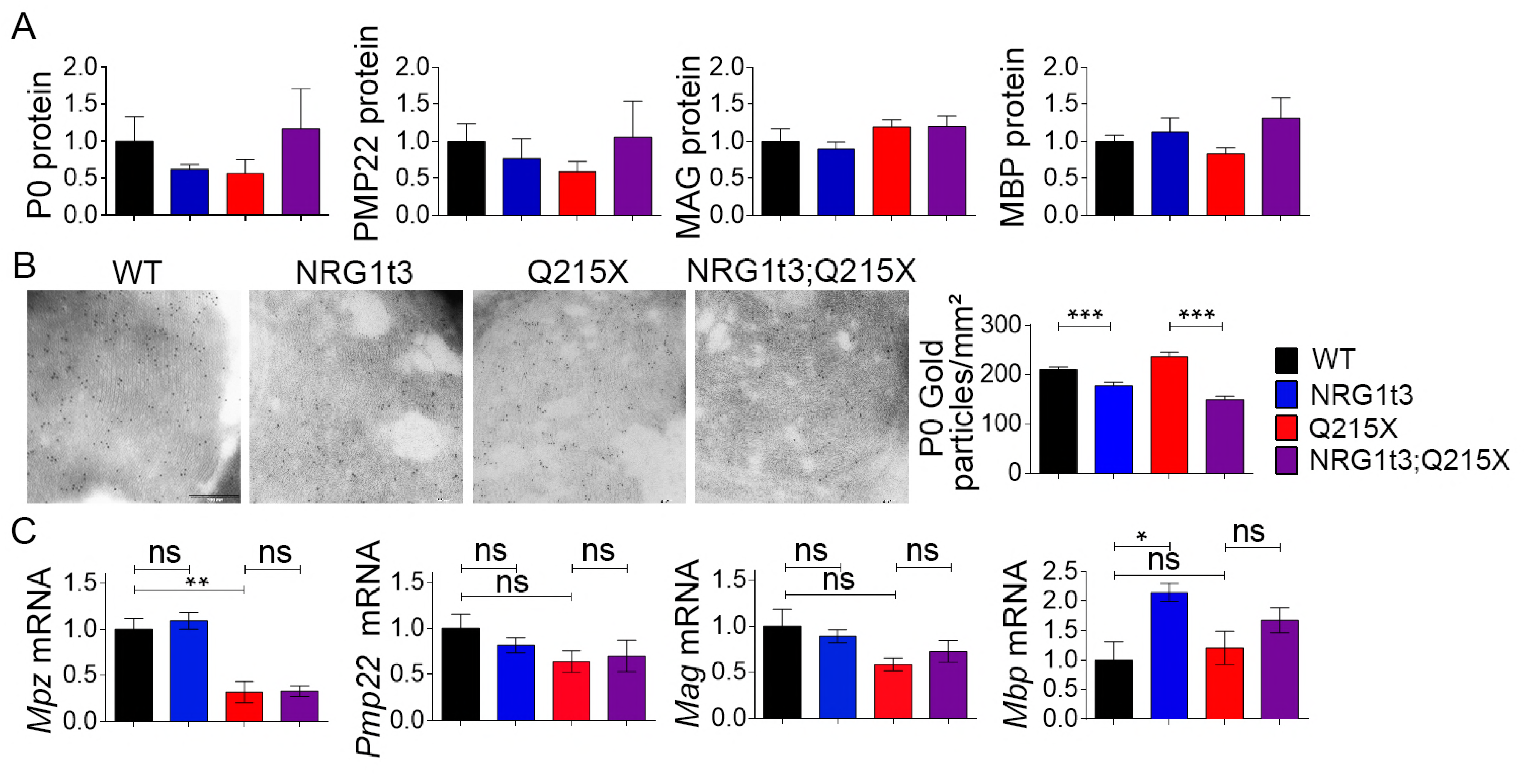
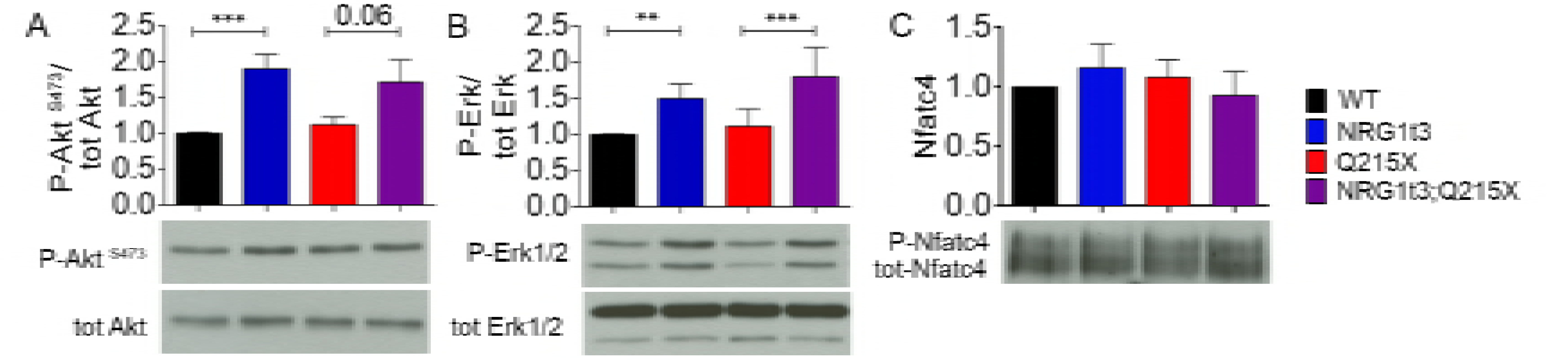
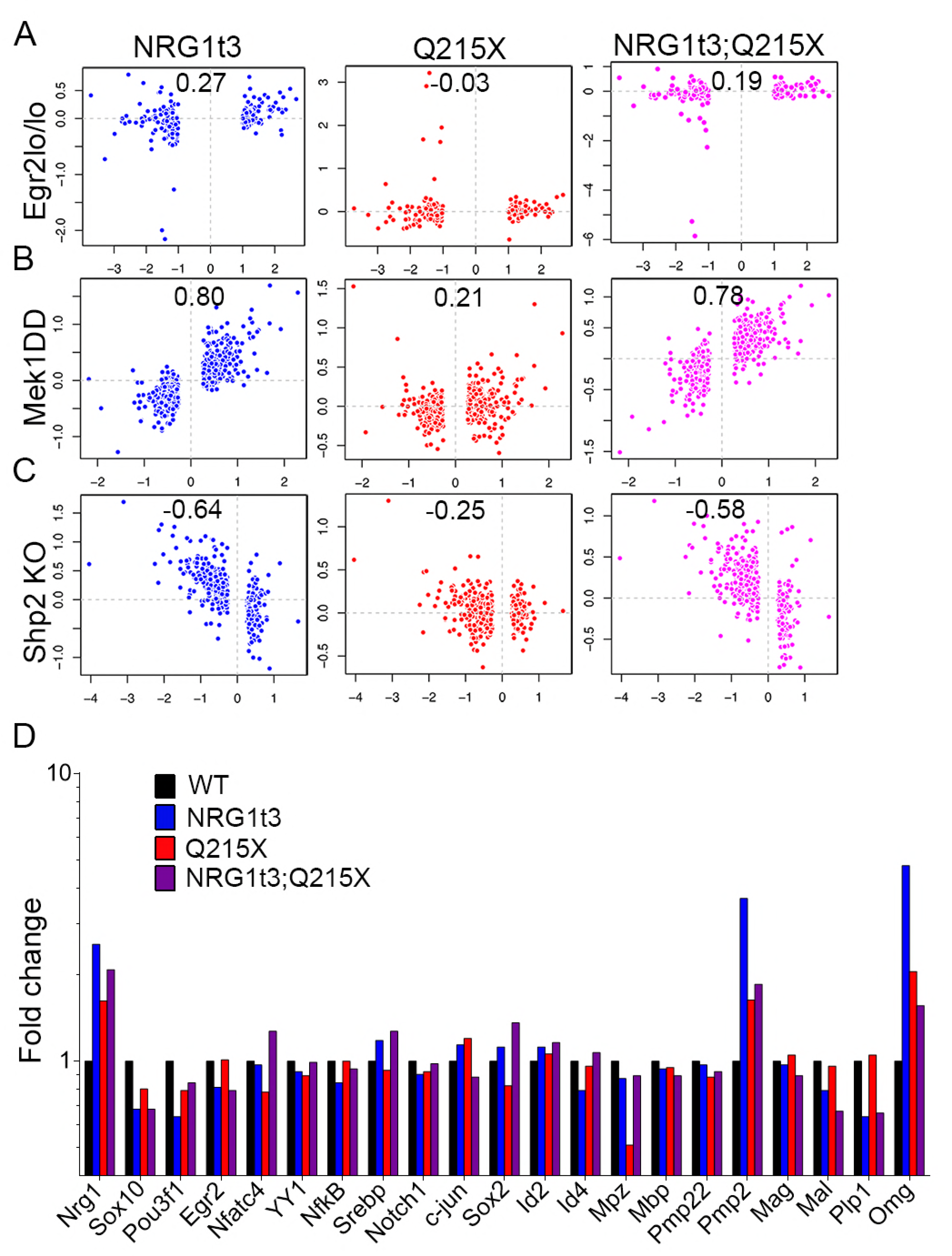
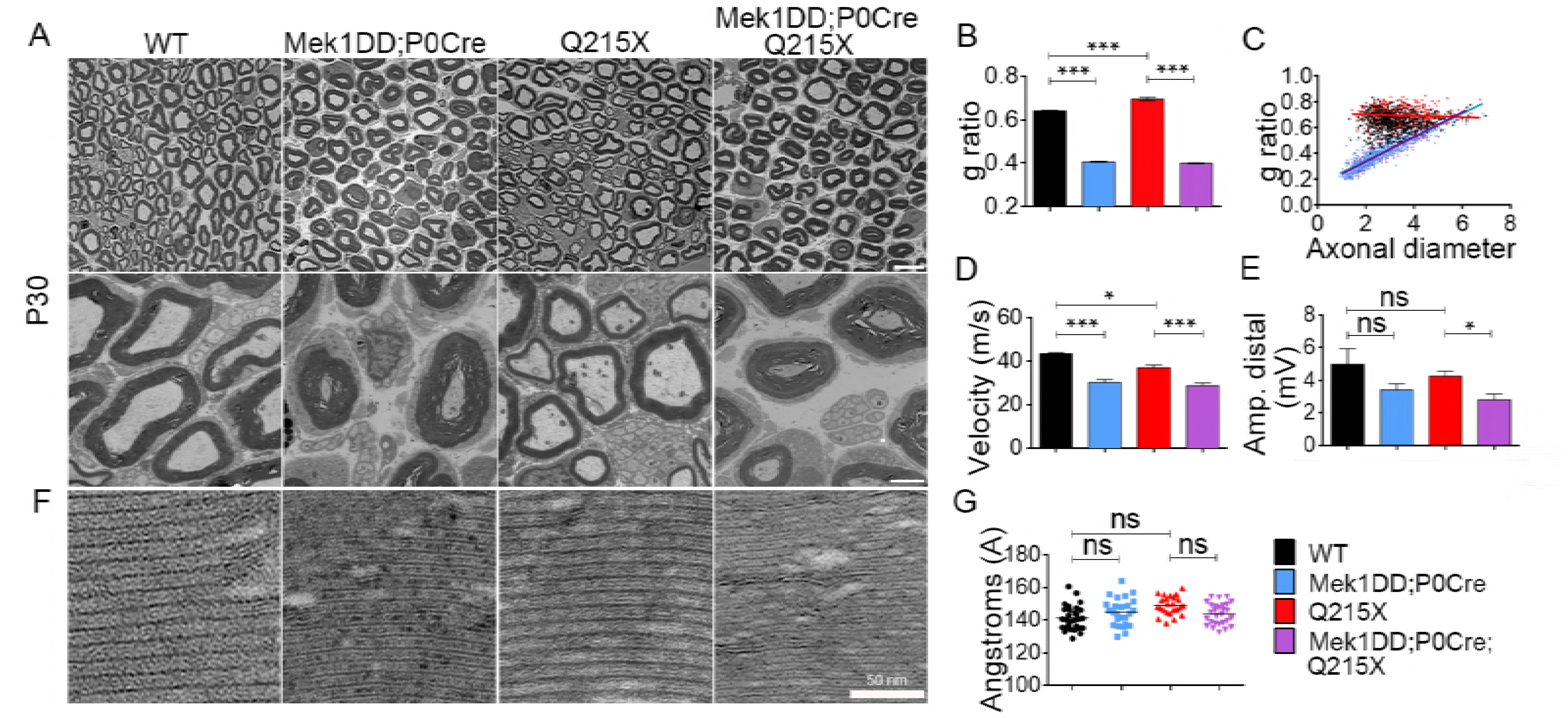
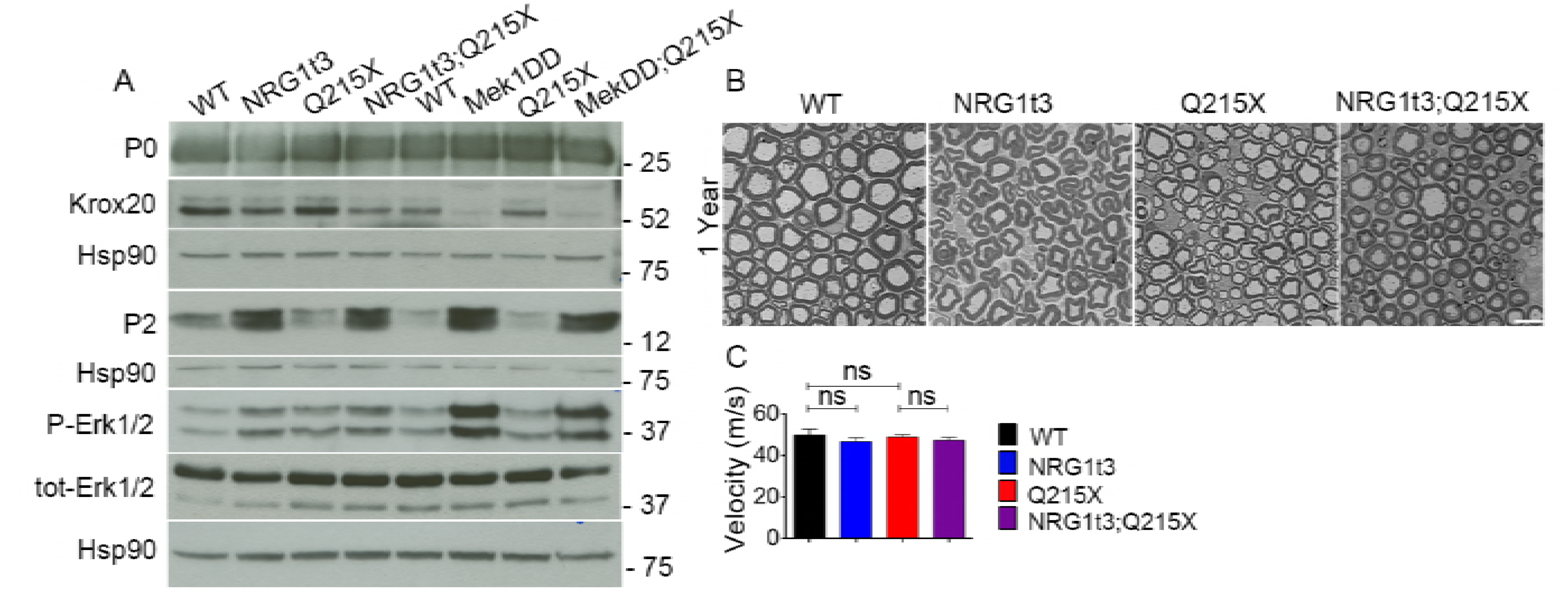
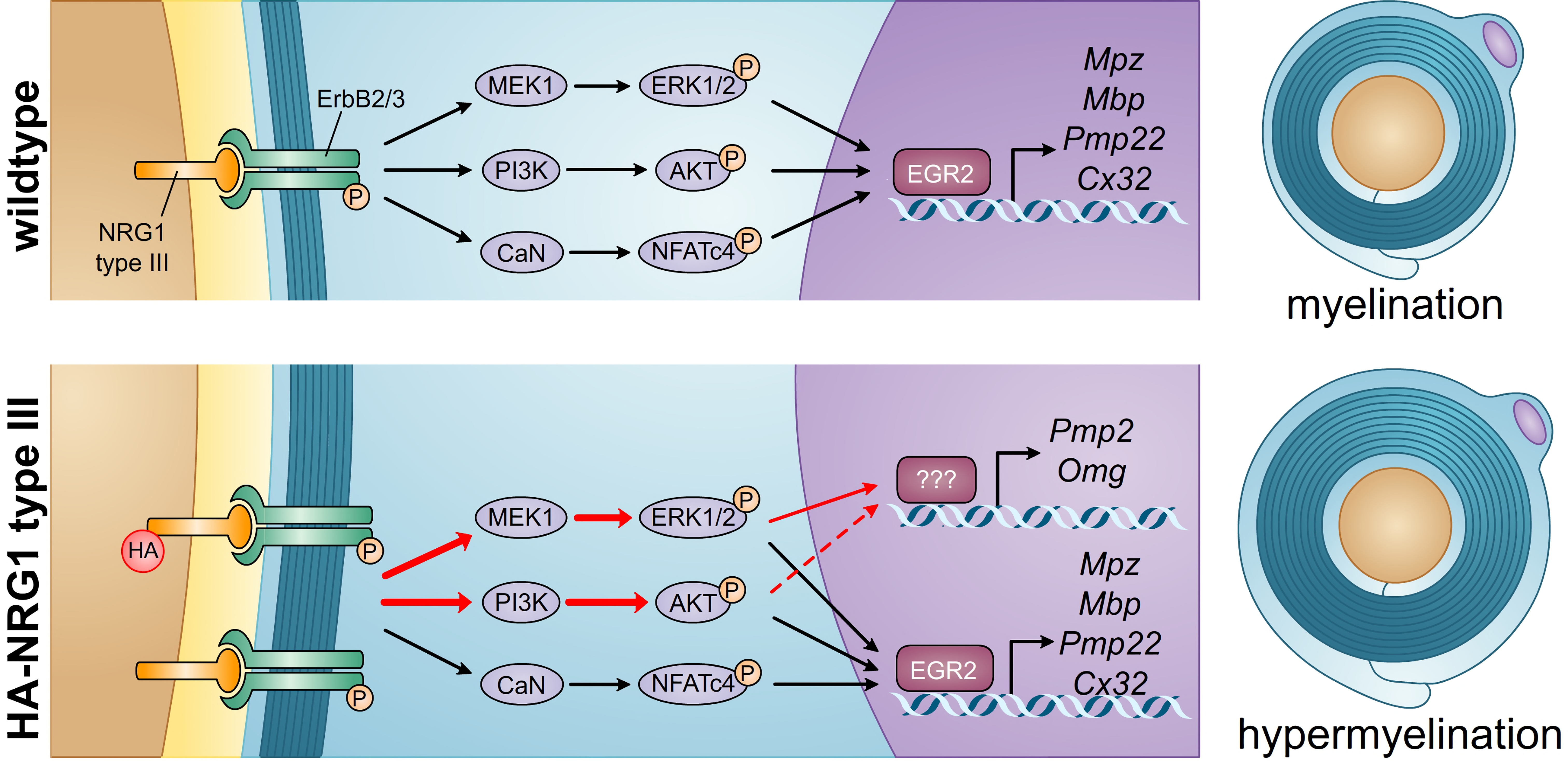
Expression of HA-NRG1 type III increases *Pmp2* mRNA and protein levels. (**A**) Relative quantification of *Pmp2* mRNA from WT, Nrg1t3, Q215X, Nrg1t3;Q215X sciatic nerves at one month. There is a significant up regulation of *Pmp2* (1.89 ± 0.16 folds in NRG1t3 and 1.81 ± 0.15 in Nrg1t3;Q215X). Error bars indicate s.e.m. Analyses were performed using three mice per genotype. Statistical analyses were performed using one-way ANOVA with Bonferroni’s multiple comparison test. *** *P* < 0.001. * *P* < 0.05. (**B**) Immunolabelling for PMP2 (in red) and P0 (in green) in cross-section from WT, Nrg1t3, Q215X, Nrg1t3;Q215X sciatic nerves at one month. PMP2 staining is increased in Nrg1t3 and Nrg1t3;Q215X sciatic nerves. Scale bar, 25μm.

## Abbreviations

CHN: congenital hypomyelinating neuropathy
HA-NRG1: hemagglutinin tagged NRG1
*Mpz*/P0: myelin protein zero
NCV: nerve conduction velocity
NRG1: neuregulin 1
*Omg*: oligodendrocyte myelin glycoprotein
P30: post-natal day 30
*Pmp2*: peripheral myelin protein 2
Q215X: Gln215 stop codon nonsense mutation

